# Long-range alteration of the physical environment mediates cooperation between *Pseudomonas aeruginosa* swarming colonies

**DOI:** 10.1101/2022.06.29.498166

**Authors:** Maxime Deforet

## Abstract

*Pseudomonas aeruginosa* makes and secretes massive amounts of rhamnolipid surfactants that enable swarming motility over biogel surfaces. But how this rhamnolipids interact with biogels to assist swarming remains unclear. Here I use a combination of optical techniques across scales and genetically-engineered strains to demonstrate that rhamnolipids can induce agar gel swelling over distances > 10,000x the body size of an individual cell. The swelling front is on the micrometric scale, and is easily visible using shadowgraphy. Rhamnolipid transport is not restricted to the surface of the gel, but occurs through the whole thickness of the plate and, consequently, the spreading dynamics depends on the local thickness. Surprisingly, rhamnolipids can cross the whole gel and induce swelling on the opposite side of a two-face Petri dish. The swelling front delimits an area where the mechanical properties of the surface properties are modified: water wets the surface more easily, which increases the motility of individual bacteria and enables collective motility. A genetically-engineered mutant unable to secrete rhamnolipids (D*rhlA*), and therefore unable to swarm, is rescued from afar with rhamnolipids produced by a remote colony. These results exemplify the remarkable capacity of bacteria to change the physical environment around them and its ecological consequences.

**Significance statement:** Living organisms have the ability to interact mechanically with their environment. *Pseudomonas aeruginosa*, a motile bacterium, can spread collectively on biogels, a behavior called swarming. Rhamnolipids, surfactant molecules *P. aeruginosa* make and secrete, are required for swarming. Here, I demonstrate rhamnolipids not only physically alter the biogel in the vicinity of the secreting cells, but also over distances much greater than the bacterial cell size, through gel swelling. This long-distance physical alteration can even rescue a remote colony which would not produce rhamnolipids. This work illustrates the remarkable ability of bacteria to change the mechanical property of the world surrounding them.

## Introduction

Bacteria have a remarkable ability to change the world around them through their collective behaviors (Papenfort & Bassler, 2016; Mavridou *et al*., 2018; Ratzke & Gore, 2018). Some of these changes are physical : bacteria interact mechanically with their environment Persat *et al*. (2015) and they are also able to feedback on the physical environment (Berk *et al*., 2012; Chew *et al*., 2014; Tropini, 2021).

Petri dishes with an agar biogel are widely used to investigate collective bacterial motility and its interplay with other biological processes (Wadhwa & Berg, 2021). This approach was used to uncover chemotactic genes (Greenfield *et al*., 2009; Colin *et al*., 2021), mechanisms of action of pharmaceutical molecules (Mirzoeva *et al*., 1997), evolutionary dynamics (Baym *et al*., 2016), quorum sensing (Daniels *et al*., 2004; Kamatkar & Shrout, 2011), and other social behaviors of bacteria (Jeckel *et al*., 2019; Badal *et al*., 2021; Monaco *et al*., 2022).

Agar gel serves as a physical and chemical substrate: cells are inoculated at the surface of the gel, they proliferate by consuming essential nutrients supplemented to the gel. In that regard, an agar plate is generally viewed as a passive element merely providing mechanical support, water, and nutrients. During colony growth, bacteria secrete osmolytes (exopolysaccharides), which draw water from the gel to equilibrate osmotic imbalance and contribute to colony swelling and expansion, in non-motile (Dilanji *et al*., 2014; Seminara *et al*., 2012) as well as in motile colonies (Ping *et al*., 2014; Yang *et al*., 2017; Rhodeland *et al*., 2020; Ma *et al*., 2021). While colony morphogenesis modeling attempts sometimes include nutrient and water depletion (Srinivasan *et al*., 2019; Yan *et al*., 2017), structural changes of agar gel are rarely considered.

In the case of the bacterium *Pseudomonas aeruginosa*, a common opportunistic pathogen, another gel modification needs to be considered: *P. aeruginosa* secretes rhamnolipids (Abdel-Mawgoud *et al*., 2010), a family of glycolipid surfactants of strong interest in medicine (Thakur *et al*., 2021) and industry (Varvaresou & Iakovou, 2015; Gudinñ *et al*., 2015), in particular for their antimicrobial properties, their capacity to emulsify oil and participate in bioremediation, and their good environmental compatibility and biodegradability. Secretion of rhamnolipids, under metabolic control (Boyle *et al*., 2015; Santamaria *et al*., 2022), allows the colony to rapidly and collectively spread into a branched shape, a phenotype called swarming (Copeland & Weibel, 2009; Kearns, 2010). Mutants that do not secrete rhamnolipids are unable to swarm (Caiazza *et al*., 2005; Xavier *et al*., 2011).

Rhamnolipids are secreted as a family of glycolipid molecules, made of one or two fatty acid chains (primarly 3-(3-hydroxyalkanoyloxy)alkanoic acid, HAA) associated with one or two rhamnose groups, called monorhamnolipids (mono-RLs) or dirhamnolipids (di-RLs) (Abdel-Mawgoud *et al*., 2010). The contribution of each molecule to swarming motility has been explored (Yeung *et al*., 2009), but the literature is inconsistent: (Caiazza *et al*., 2005) reported HAA is merely is wetting agent and a wild-type colony growing on a HAA-supplemented plate swarms normally. In contrast, (Tremblay *et al*., 2007) showed HAA is a strong repellent for swarming colonies and a wild-type colony growing on a HAA-supplemented plate is inhibited. A recent report from the same group (Morin & Déziel, 2021) showed that mutants producing only HAA are still able to swarm and form branches. These conflicting data call for a deeper understanding of the mechanisms underlying rhamnolipids-assisted swarming motility, in particular about the exact contribution of HAA, mono-rhamnolipids, and di-rhamnolipids. In the literature, two main mechanisms were suggested: (i) rhamnolipids increase wettability of the agar surface, (ii) they create gradients of surface tension that cause the colony to move outwards (Marangoni effect).

Due to their surfactant nature (Yang *et al*., 2021), rhamnolipids are often said to *lubricate the surface* (Boyle *et al*., 2015), decrease *friction against the gel* (Hölscher & Kovács, 2017), *act as a wetting agent* (Tremblay *et al*., 2007), or *lower surface tension* (Yang *et al*., 2017; Rütschlin & Böttcher, 2020). Addition of synthetic surfactants to a swarming plate is known to greatly improve colony spreading dynamics (Pamp & Tolker-Nielsen, 2007; Yang *et al*., 2017). Change of wettability, induced by biosurfactants, can even enable flow of bacterial suspension along solid surfaces and through unsaturated porous material (Yang *et al*., 2021). It is clear the molecular details of the gel and in particular the gel surface are crucial factors in how the colony behaves: patterns and spreading rate depend on the hardness of the gel (via agar concentration) (Kamatkar & Shrout, 2011; Mattingly *et al*., 2018), the method of gel preparation (Tremblay & Déziel, 2008), but also the choice of gelling agent (Morin & Déziel, 2021), as well as any modification of its viscoelastic properties with substances like mucin or carboxymethyl cellulose (Yeung *et al*., 2012), or polyethylene oxide (Yang *et al*., 2017).

Early works reported how surfactant-assisted spreading of abiotic liquid film could yield to branching. This has been explored on liquid and solid substrates (Matar & Troian, 1999; Matar & Craster, 2009), and was extended to the case of *P. aeruginosa* swarming colonies (Du *et al*., 2011, 2012; Fauvart *et al*., 2012; Trinschek *et al*., 2018). In those works, rhamnolipids were modeled as a thin film of insoluble surfactants secreted by the colony and diffusing at the surface of a gel. Differences of surfactant concentration inside and outside the colony yield to gradients of surface tension and induced a branching instability (a mechanism known as Marangoni effect). These modeling efforts ignored what was happening on the agar side, where a precursor film was sometimes observed. In particular, rhamnolipids are soluble in water (Abdel-Mawgoud *et al*., 2009) and these molecules are substantially smaller than the agar gel pores. Therefore, they are expected to diffuse though the entire agar gel, a hydrogel, whose structure can be modified through swelling (Bibi *et al*., 2019; Li *et al*., 2021).

Last, the expansion of a *P. aeruginosa* swarming colony can be divided into two components: spreading outwards and branching. Different theories attempt to explain these distinct processes. It is believed that the outward spread is due to an osmotic influx of water and a rhamnolipid-induced increase in surface wettability. Branching, which was recently found to optimize colony growth (Luo *et al*., 2021), may be caused by a rhamnolipid-induced Marangoni effect, or alternatively by diffusion-limited growth or chemotaxis (Deng *et al*., 2014; Giverso *et al*., 2015). In this context, the knowledge of the contribution of each rhamnolipid congeners (HAA, monorhamnolipids, di-rhamnolipids) to each mechanism in still very limited.

Here, I use a combination of imaging methods across scales to demonstrate that rhamnolipids secreted by the colony alter the physical properties of the agar gel, not only locally where they are secreted, but also all more widely around the bacteria. Moreover, their transport is not restricted to the surface. I find, instead, that the rhamnolipids imbibe the whole gel, which yields to gel swelling with a sharp swelling front. Bulk transport of rhamnolipids directly affects the expansion rate of the region imbibed by rhamnolipids. Gel imbibition by rhamnolipids cover so large distances that rhamnolipid-deficient colonies can be remotely rescued. More generally, this is an example of the remarkable ability of bacteria to change the mechanical properties of the world surrounding them.

## Experimental Procedures

### Bacterial strains and growth conditions

The bacterial strain (*P. aeruginosa* laboratory strain PA14) and its mutants used in this study are described in Table 1. Bacterial cells were routinely grown in LB at 37ºC with aeration. Swarming medium (2.37 M Na2HPO4, 1.81 M KH2PO4, 4.67 M NaCl, 1 mM MgSO4, 0.1 mM CaCl2, 5 g/L casamino acids (Bacto, BD)) was solidified with agar (Bacto, BD), following previously described recipes (Xavier *et al*., 2011; Deforet *et al*., 2014). Agar concentration, unless specified otherwise, is 0.5 % (w/v). Overnight culture of bacteria were washed twice in PBS and 2 µL of the washed suspension were used to inoculate a swarming plate in the center. The plates were then flipped and placed inside a 37ºC microbiological incubator. D*rhlA:P*_*BAD*_*rhlAB* colonies were grown on 1% L-arabinose swarming plates to induce expression of the *rhlA* gene and robust secretion of rhamnolipids.

**Table 1:**
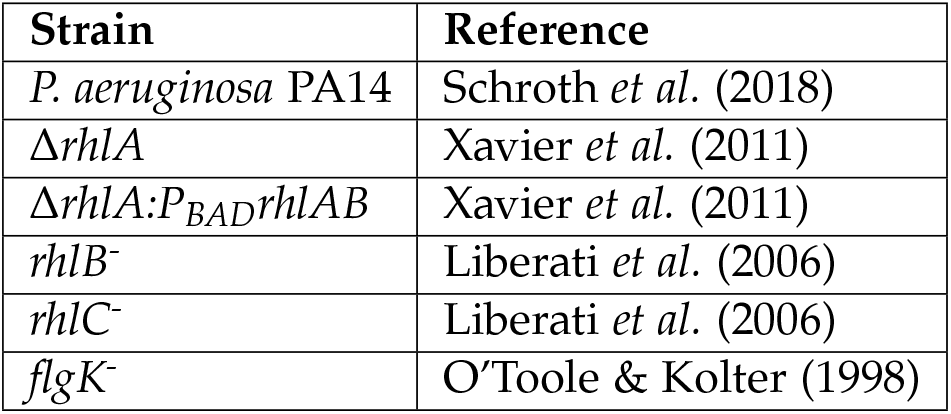
Strains used in this study

### Whole swarming colony

An automated device was built inside a microbiological incubator (Heratherm IGS 100, ThermoFisher Scientific), for fluorescence, shadowgraphy, and brighfield imaging of a 10-cm diameter Petri dish. A LED light source (pE-4000, Coolled, UK) was used for fluorescence excitation. A set of lenses and mirrors were used to bring an even illumination pattern onto the plate. A single LED light source (Thorlabs, USA) was positioned on one side of the chamber for shadowgraphy imaging (see Figure S1 for a schematic of the imaging device). Generic white LED strings (Mouser, France) were positioned above the plate for brightfield imaging. Images were acquired with a CMOS camera (Cellcam Centro, Cairn Research, UK) with a macro lens (Navitar MVL7000, Thorlabs, USA). A dual-band emission filter (59010m, Chroma, USA) was mounted between the camera and the lens, for green and red fluorescence imaging. All devices were controlled with µManager (http://www.micro-manager.org).

### Sessile droplets

Plates and swarming colonies were prepared as explained above. Plates were taken out of the incubator and placed under a Axiozoom V16 macroscope (Zeiss), set at the lowest magnification (field of view is 25.7×21.5mm), equipped with a CMOS camera (BlackFly S, FLIR, USA), and illuminated through a transilluminator (Zeiss) with a mirror position that mimics DIC illumination. 1 µL droplets of swarming media (recipe identical to that of the swarming plate, but without agar) were deposited on the surface of the gel, either inside or outside the swelling front generated by the colony. A movie was recorded as tens of droplets were deposited. Footprint diameters were measured, immediately after deposition, using ImageJ.

### Fluorescent bead tracking

For quantifying gel swelling, 1 µm fluorescent polysterene beads (F13083, ThermoFisher Scientific) were added during the swarming plate preparation (final concentration of 10^*5*^ *beads/mL)*. Colony were grown inside a microscopy incubator (Oko-Lab, Italy), mounted an a IX-81 Olympus inverted microscope, equipped with a LED light source (pE-4000, Coolled, UK). Images were taken with a 20x objective and a CMOS camera (BlackFly S, FLIR, USA), every minute, for 30 minutes, while the swelling front was passing through the field of view (confirmed by phasecontrast imaging). Diffraction pattern detection, beads detection, tracking, and calculation of the vertical displacement were performed with custom-made routines in MATLAB (MathWorks, USA). Calibration between diffraction pattern radius and vertical position was done by taking a vertical stack of images.

### Profilometry

Height profiles were measured with a Zegage Pro interferometry profilometer (Zygo) equipped with a 5x Michelson objective (Nikon). An acquisition took 5 seconds (for one field of view 2×2.7 mm), and 20 positions were stitched together to reconstruct the whole height profile (Figure 1E). For Figure 5C, the profiles were measured when the diameter of the rhamnolipids-imbibed area was approximately 3 cm in diameter. Four positions were recorded per plate (the 4th roots of unity). Each profile was then measured perpendicularly to the swelling front. The *X* = 0 location was identified as the inflection point of the profile. The reference height *Z* = 0 was imposed at location *X* = *-*500 µm and *X* = *-*400 µm, which tilt-corrected the whole profile. Heights in Figure 5C were measured at *X* = 1000 µm. For Figure S5, a 3×3 stitch acquisition was performed 5 minutes after the liquid was entirely absorbed by the gel.

**Figure 1:**
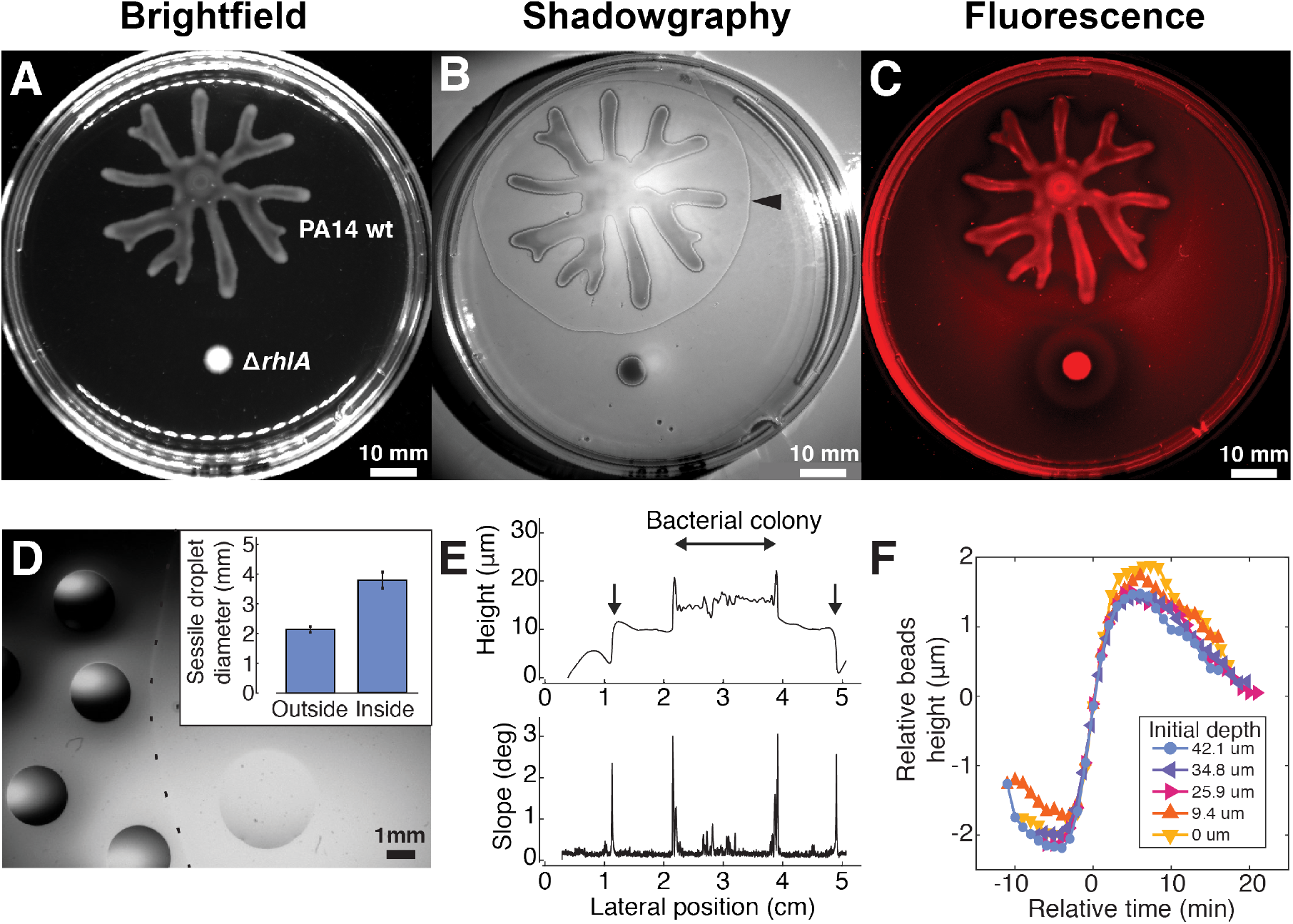
Rhamnolipids induce agar gel swelling. A-B-C: Wild-type *P. aeruginosa* and D*rhlA* colonies were grown on an agar gel. The former swarmed out and formed a branched colony, whereas the latter did not swarm. A: Brightfield image. B: Shadowgraphy image reveals a darker line (depicted by a black arrowhead) that corresponds to a swelling front. C: Fluorescence image, for Nile Red imaging, confirms the darker line correponds to a boundary of an area imbibed by rhamnolipids (see Figure S2 for a montage combining panels B and C). D: An illustrative micrograph showing the sessile droplets experiments. Dashed line depicts the swelling front. The source colony is on the right side. Inset: comparison of the wetting footprint radius in the two conditions. Error bars are standard deviation. E: Top: a height profile of a gel, obtained from optical interferometry. The downward arrows indicate the swelling front. Bottom: The local slope of the height profile. F: Vertical positions of fluorescent microbeads reveals gel swelling. The time axis for each trajectory is shifted using time synchronization from lateral displacement data (See Figure S3). Colors code for initial depth of each bead.

### Stepwise substrate

A 1.5 mm thick sheet of polydimethylsiloxane (PDMS) was made following standard protocol (RTV615A+B, 10:1, Momentive Performance Materials). A 1×2 cm slab was cut out of the PDMS sheet and was placed in a Petri dish. 20 mL of agar gel was then poured on top of the substrate, following usual swarming plate protocol.

### Two-face Petri dish

The bottom of a Petri dish was drilled with a 10-mm drill bit. Holes were deburred with a 13-mm drill bit. Dishes were sterilized in 70% ethanol. The bottom side of each hole was taped with a piece of Parafilm, which was peeled off after the agar gel was poured and solidified.

### Single-cell motility

A suspension of D*rhlA* cells growing in liquid swarming media was spun down. A 20 µL plastic pipette tip was used to sample cells from the pellet and to transfer cells onto the gel surface (approximately 2 mm from the swelling front identified in shadowgraphy). One second videos (46 frames-per-second) were acquired every 5 minutes, at 10x magnification, in phase contrast, with an IX-81 Olympus inverted microscope. Auto-focus was performed before each acquisition. Image analysis was performed independently for each video: Local density was measured on the first frame of the video, by thresholding the phase-contrast image, followed by local averaging. The Density Index, defined to be between 0 (no cell) and 1 (cells form a uniformly dark population) is therefore the proportion of neighboring pixels that contains cells. To evaluate motility, the difference between the maximum projection and the minimum projection across the time-stack yields to a map where pixels were bright if their grey values changed during the video. Then, a threshold was applied and the result was locally average to produce a map of Speed Index between 0 (no pixel change) and 1 (all pixels have change values). This coarse-grained quantification is valid as long as cells are not too dense and cells do not move too much during one video (therefore, the analysis was limited to the region of the field of view where cells did not pile up and only one-second long videos were considered). To check the robustness of the method, various thresholding values have been tried, without significantly affecting the results.

## Results

### Rhamnolipids cause biogel swelling

Wild-type *P. aeruginosa* colony spreads on soft agar by forming a branched colony, a behavior called swarming (Kearns, 2010). *rhlA*, a gene involved in production of rhamnolipids, is required for swarming: a D*rhlA* colony, which does not produce rhamnolipids, is unable to swarm (Figure 1A). The presence of rhamnolipids around a colony has been revealed using methylene blue (Yeung *et al*., 2009; Abdel-Mawgoud *et al*., 2010). Shadowgraphy, a non-destructive, optical method that reveals non-uniformities in transparent media by casting a shadow onto a white background (Settles, 2001) (see Figure S1 for a schematic of the imaging device), confirmed that an area of the gel around the rhamnolipid-producing colony was modified: a thin darker line surrounded the colony (Figure 1B). Timelapse imaging confirmed this optical modification originated from the colony and propagates outward (See Movie S1). Since no modification emerged from a D*rhlA* colony, I hypothesized variation of the optical properties of the gel were due to secretion of rhamnolipids.

Since rhamnolipids and some of their precursors are lipids, they could be localized using a lipid dye. Nile Red, whose emission spectrum depends on the polarity of the solvent (Teo *et al*., 2021), was mixed to the agar gel during gel preparation (Morris *et al*., 2011). Using fluorescence imaging to analyze solvent polarity around colonies, there was a distinct difference between wildtype and D*rhlA* colonies, with a sharp variation coinciding with the darker line that surrounds the wild-type colony (Figure 1C and S2). This provides even stronger evidence that the darker line marks the rhamnolipid range. I also measured the wetting property of the gel surface by performing sessile droplets experiments (Banaha *et al*., 2009; Santamaria *et al*., 2022). Water droplets were deposited on each side of the darker line localized by shadowgraphy. On average, the droplet footprint radius was 3.78 *±* 0.27 (SD) mm on the colony side, and 2.13 *±* 0.09 (SD) mm on the other side (Figure 1D). Neglecting gravity and therefore assuming a spherical cap shape allow the calculation of contact angles of 22.7º and 35.6º, respectively. The wetting footprint size was found to be uniform on the colony side, independent of the distance to the colony or the distance to the darker line. Similarly, the wetting footprint size was found to be uniform also outside the darker line. This confirms that the darker line marks a sharp limit of the rhamnolipid range, with surface properties that correlates with the presence of rhamnolipids. (Of note, the resolution of the sessile droplet experiment does not enable to measure the spatial distribution of rhamnolipid concentration, which was previously performed using LC/MS (Tremblay *et al*., 2007); the data merely demonstrate a contrast of concentration inside/outside the darker line.)

To understand the origin of the darker line visible in shadowgraphy, I had to first evaluate its 3D profile. Using an optical profilometer (a device capable to measure 3D profile of a reflective surface with a nanometric resolution), I measured a jump of 8 µm, with a maximal slope of 3º (Figure 1E). The jump could emerge from two possible mechanisms: either it corresponds to a thick layer of rhamnolipid molecules covering the surface of the agar gel, as assumed by previous works (Fauvart *et al*., 2012; Trinschek *et al*., 2018), or alternatively rhamnolipids infiltrate the gel and the jump corresponds to a gel swelling front. I embedded micrometric fluorescent beads during preparation of the gel and performed 3D-tracking to follow each bead while the line passed through the field of view. If rhamnolipids cover the gel surface, beads should not move. If rhamnolipids diffuse through the gel and make it swell, beads should move up. Data shown in Figure 1F confirms a vertical movement of all the beads, synchronized with the passing line (see also Movie S2). The swelling amplitude was uniform across the first 50 µm (optical limitations of fluorescence microscopy hamper deeper observations). Of note, beads also moved laterally with a similar amplitude: they first moved outwards and then inwards (this corresponds to a dilatation wave, see Figure S3 for more details). This confirms rhamnolipids make the agar gel swell over large distances, and the darker line corresponds to a swelling front that is so steep that it can be seen by a naked eye (Xavier *et al*., 2011) and by shadowgraphy.

Shadowgraphy enables to locate the boundary of the rhamnolipid range on the plate, but it is unable to measure the spatial distribution of rhamnolipid congeners within the detected range. Yet, combining rhamnolipid mutants with shadowgraphy observations reveals information about the timing of secretion of various rhamnolipid congeners. I located the position of the swelling front with respect of time and this displayed two phases (Figure S4A): for the first 6 hours, a swelling front emerged from the colony, traveled for 10 mm, and stalled. Then it regained speed and kept on traveling across the entire plate. I compared this evolution with that of two transposon mutants: the *rhlB*^*-*^ *mutant is u*nable to convert HAA into mono-rhamnolipids, and the *rhlC*^*-*^ *mutant is u*nable to convert mono-rhamnolipids into di-rhamnolipids (Abdel-Mawgoud *et al*., 2010) (Figure S4B). The two phases were identified for the *rhlC*^*-*^ *plate too*. In contrast, only the first phase was identified for the *rhlB*^*-*^ *plate, indi*cating the second phase was enabled by mono-rhamnolipids. Since a D*rhlA* colony, unable to convert b-hydroxylacyl-ACP into HAA, does not generate any swelling front at all, I can estimate that the first phase is dominated by HAA (see the table in Figure S4D for a summary of these findings). In the rest of this work, all studied swelling fronts correspond to the second phase.

Finally, since mono- and di-rhamnolipids and HAA are surfactants (Deziel *et al*., 2003; Abdel-Mawgoud *et al*., 2010), I tested if their ability to swell agar gels was a generic feature of surfactant molecules. I tested Triton X-100, Tween 20, and Tween 80, nonionic surfactants, and SDS (Sodium dodecyl sulfate), a anionic surfactant. 3D profilometry and shadowgraphy confirmed that agar gel was swollen by synthetic surfactants, as well as the supernatant of the liquid culture of wildtype *P. aeruginosa* and *rhlC*^*-*^, *but not a*fter exposure to D*rhlA* and *rhlB*^*-*^ *supernatant* (Figure S5).

### Rhamnolipids imbibe the bulk of the gel, not the surface

If transport of rhamnolipids is restricted to the gel surface, or if rhamnolipids diffuse through the gel following normal diffusion, their spreading rate should not depend on the local thickness of the gel. On the contrary, if Darcy’s law governs rhamnolipids’ transport, their spreading dynamics should depend on the local section of the gel. I used gels with varying thicknesses to test this hypothesis and check whether rhamnolipids imbibed the whole gel or only the upper part. First, I used gels with stepwise thickness. I placed a thick inert obstacle at the bottom surface of the Petri dish and poured the agar gel on top of them. The thickness of obstacles (1.5 mm) was chosen to be smaller than the gel thickness (3.5 mm) to make sure the upper surface remained flat. To simplify the geometry of the system and keep a local source of rhamnolipids, I inoculated a non-motile mutant (*flgK*^*-*^*) along a l*ine. The swelling front coming out of the colony was found to propagate faster on top of obstacles (Figure 2A), where the gel section was smaller, and slower on top of thicker sections of the gel, which is compatible with a transport of rhamnolipids through the whole gel. This effect was also found in gels of various thicknesses. Once again, thinner gels induced faster traveling speed of the swelling front (Figure 2B), in agreement with transport through the gel. Intriguingly, while thinner gels induce faster spreading of the swelling front, they also induce smaller swarming colonies (Figure S6). Finally, I tested the swelling speed in a gel with gradual change of thickness. Pouring liquid agar on a tilted dish resulted in a gel with a spatial gradient of thickness: on this substrate, the swelling front advanced faster on the thinning side, slower on the thickening side, and at intermediate speed on the transverse directions (Figure 2B). Colony did not swarm at all on plates lower than 7 mL. They gradually were larger and larger on thicker gels and the colony size was maximal on plates of the nominal volume (20 mL).

**Figure 2:**
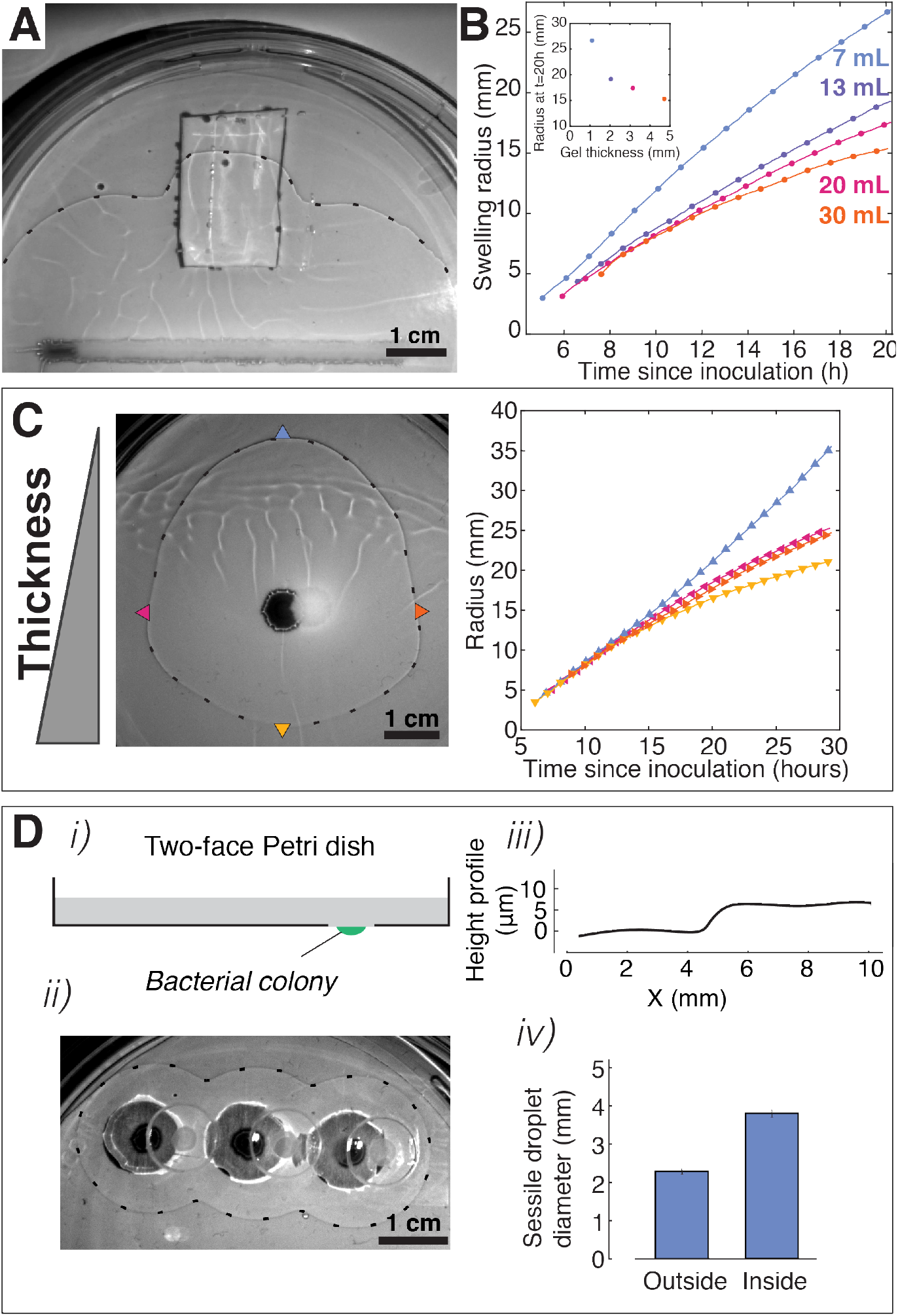
A: Snapshot of the swelling front on a plate with an obstacle. The non-motile colony, inoculated along a line, is visible near the bottom of the image. B: Positions of the swelling front as a function of time, for four plates with different thicknesses of gel. C: Left: Snapshot of the swelling front advancing on a tilted gel. Right: Positions of the swelling front as a function of time, for 4 locations depicted with arrowheads (top, bottom, left, and right). D: Two-face Petri dish confirms rhamnolipids spread through the gel. i) Schematic of a two-face Petri dish. ii) Shadowgraph of the swelling front emerging from three colonies, located on the other side of the gel. iii) Height profile obtained from optical profilometry. iv) Sessile droplet experiment confirms a significant difference in wetting property on either side of the swelling front. Error bars are standard deviation.

To further demonstrate transport of rhamnolipids through gels, I designed a “two-face” Petri dish. I inoculated bacteria on the lower side of the dish and a swelling front was observed on the upper side. The upper surface was characterized with shadowgraphy, profilometry, and sessile droplets (Figure 2D). These measurements confirmed that a rhamnolipid-induced swelling front propagated to the upper surface, even though rhamnolipids were secreted from the lower surface.

### Rhamnolipids enable surface motility at the single-cell level

Rhamnolipids, transported within the gel, are required for the colony to spread at the surface of the gel. To better understand the interplay between bulk transport of rhamnolipids and surface motility, I used D*rhlA* mutant, which does not secrete rhamnolipids but is motile, as a sensor of ability to move on the gel as a function of presence of rhamnolipids. A wild-type colony was grown on agar gel and a small amount of planktonic D*rhlA* cells were seeded on the naked gel. Individual cells were found to be initially mostly non-motile. Motility was detected only in denser areas. The passage of the swelling front dramatically perturbed spatial organization of the cells (Figure 3A and Movie S3), and hampered direct comparison before/after. In particular, following individual cells was impossible. Instead, motility was assessed by coarse-grained image analysis (see Methods). The motility-density dependence was quantified and plotted against time (Figure 3B-C). Even though cells gradually became more motile with time, incoming rhamnolipids substantially increased cell motility, at all densities (Movie S4). Therefore, rhamnolipids are used by bacteria to alter their physical environment over large distances and enable surface motility.

**Figure 3:**
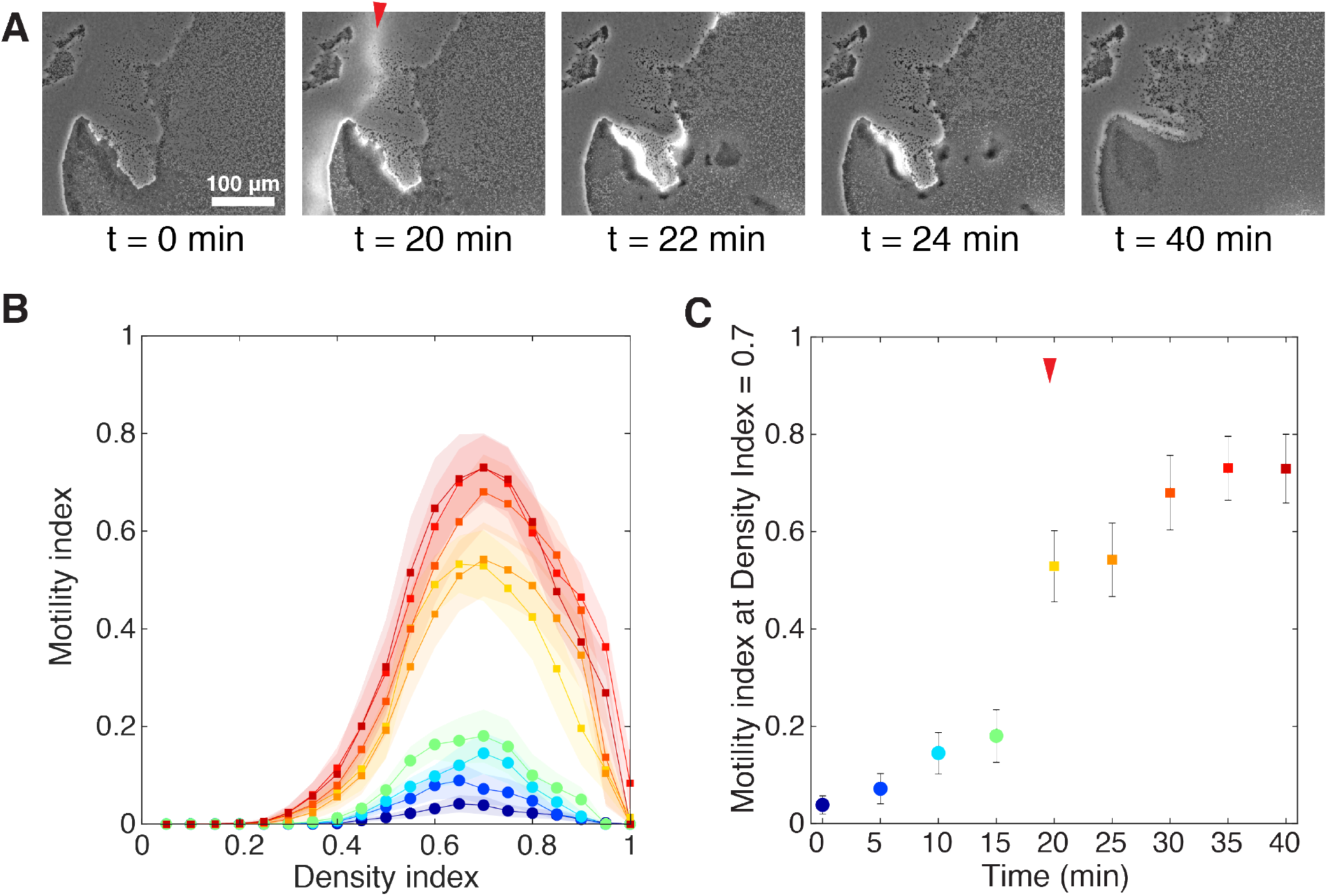
A: Snapshots of micrographs showing D*rhlA* cells before and after the swelling front passed through. A red arrowhead depicts the swelling front, visible as a white halo in phase contrast imaging, and moving from the left to the right of the field of view. B: Motility-density analysis reveals rhamnolipids greatly facilitate single-cell motility. Color code is defined in panel C (one curve every 5 minutes). Shading represents standard deviations. C: The effect of rhamnolipids on motility is abrupt, and corresponds to the passage of the swelling front (depicted by the red arrowhead). Error bars represent standard deviations.

### Swarming rescue experiments confirm rhamnolipids diffuse across the biogel

This microscopic-scale rescue of motility D*rhlA* mutants inspired me to test if such distant rescue could occur at a macroscopic scale. A plate was inoculated with wild-type cells on one side, and with D*rhlA* cells on the other side. As already showed in Figure 1, the wild-type colony secretes rhamnolipids (inducing a swelling front) and swarms into a branched shape. The D*rhlA* colony does not secrete rhamnolipids and does not swarm. However, wild-type colony spreading closely follows the swelling front. To minimize the risk of the secreting colony coming into contact with the D*rhlA* colony, I turned into the D*rhlA:P*_*BAD*_*rhlAB* mutant, whose rhamnolipid promoter expression is induced by a chemical cue, L-arabinose. The D*rhlA:P*_*BAD*_*rhlAB* colony dedicates a large part of its metabolism to rhamnolipid secretion rather than cell proliferation (de Vargas Roditi *et al*., 2013), which yields to a swelling front traveling substantially faster than the spreading of the colony (Figure S7). As shown on Figure 4A (and Movie S5), D*rhlA* colonies started swarming a few hours after the swelling front reached it. Interestingly, the D*rhlA* colonies did not spread radially, but formed tendrils, whose size is comparable to wild-type colony branches. These tendrils swarmed outward, following the direction of rhamnolipids propagation. These observations suggest the process of branching is the same whether the colony secretes its own supply of rhamnolipids or has an outside source.

**Figure 4:**
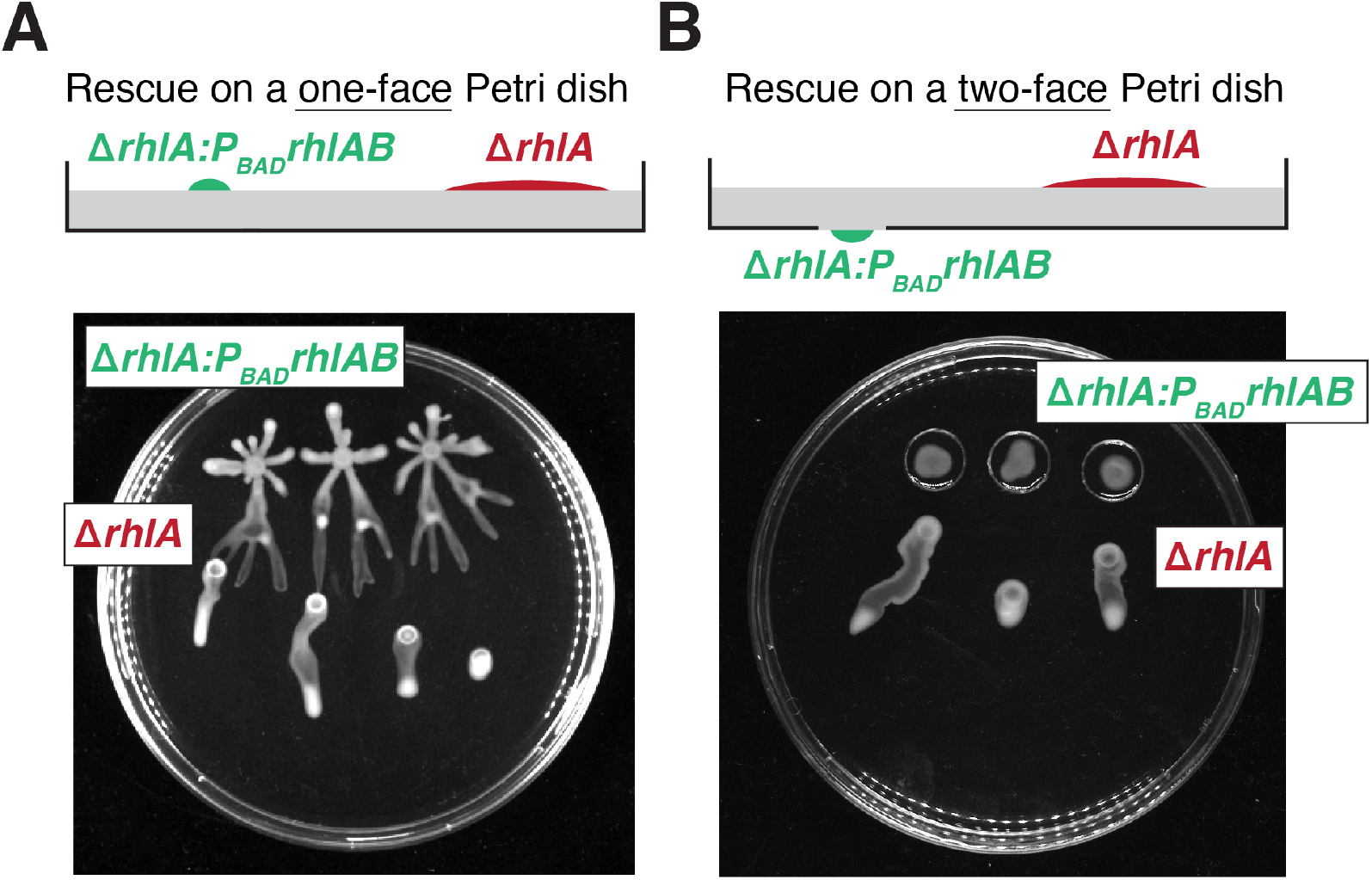
Swarming rescue experiments. A: On one-face Petri dish, D*rhlA:P*_*BAD*_*rhlAB* and D*rhlA* colonies are grown on the same side. B: On two-face Petri dish, D*rhlA:P*_*BAD*_*rhlAB* are grown on one side (within the small hole) and D*rhlA* are grown on the opposite side. Control experiment results (with D*rhlA* instead of D*rhlA:P*_*BAD*_*rhlAB*) are provided in Supplementary Figure S8. Snapshots are taken 18 hours after inoculation. Timelapse videos of the rescue experiments are available in Movie S5.

I reproduced the rescue experiment on a two-face Petri dish, with the over-producing colony (D*rhlA:P*_*BAD*_*rhlAB*) on one side and D*rhlA* on the other side (Figure 4B, Movie S5). Here again, the D*rhlA* colonies did not spread radially, but formed a tendril spreading in the direction of rhamnolipid propagation. This results confirms rhamnolipids imbibe the whole gel, physically alter it over large distance, and enable a branching process similarly on both side of the two-face Petri dish.

### Physical alteration of the gel by rhamnolipids correlates with colony shape

Since *P. aeruginosa* is capable to alter physical properties of its environment over long distances, I wanted to verify whether this would hold true for a range of environments. I used agar gels of various agar concentrations (0.4%, 0.5%, 0.6%, and 0.7%) to simulate environments of different physical properties. Agar gel elastic modulus and pore size were previously characterized and they were found to be highly dependent on agar concentration. From the literature, pore size varies from 400 nm at 0.4% to 100 nm at 0.7% (Narayanan *et al*., 2006; Cuccia *et al*., 2020); elastic modulus varies from 4 kPa at 0.4% to 18 kPa at 0.7% (Guenet & Rochas, 2006; Mao *et al*., 2016). (Most studies focused on agarose, a polysaccharide isolated from agar, but with comparable mechanical properties; rheological measurements are difficult to perform on these soft gels, and those figures must be taken with a grain of salt.)

I grew *P. aeruginosa* on plates at the four agar concentrations and confirmed a result already reported (Tremblay & Déziel, 2008; Kamatkar & Shrout, 2011; Mattingly *et al*., 2018): swarming colony morphogenesis is highly dependent on the gel properties (Figure 5A). At low concentration (0.4%), the colony was smaller and more circular than the branch colony formed at 0.5%. At 0.6%, the colony was still branched but smaller than 0.5% colony. The 0.7% colony did not spread at all. Following the same trend, macroscopic swarming rescue on two-face Petri dished occurred similarly at 0.4% and 0.5%, little spreading was visible at 0.6%, and no spreading at all at 0.7% (Figure 5B). In parallel, I characterized the gel and its alteration by secreted rhamnolipids, for the four agar concentrations.

**Figure 5:**
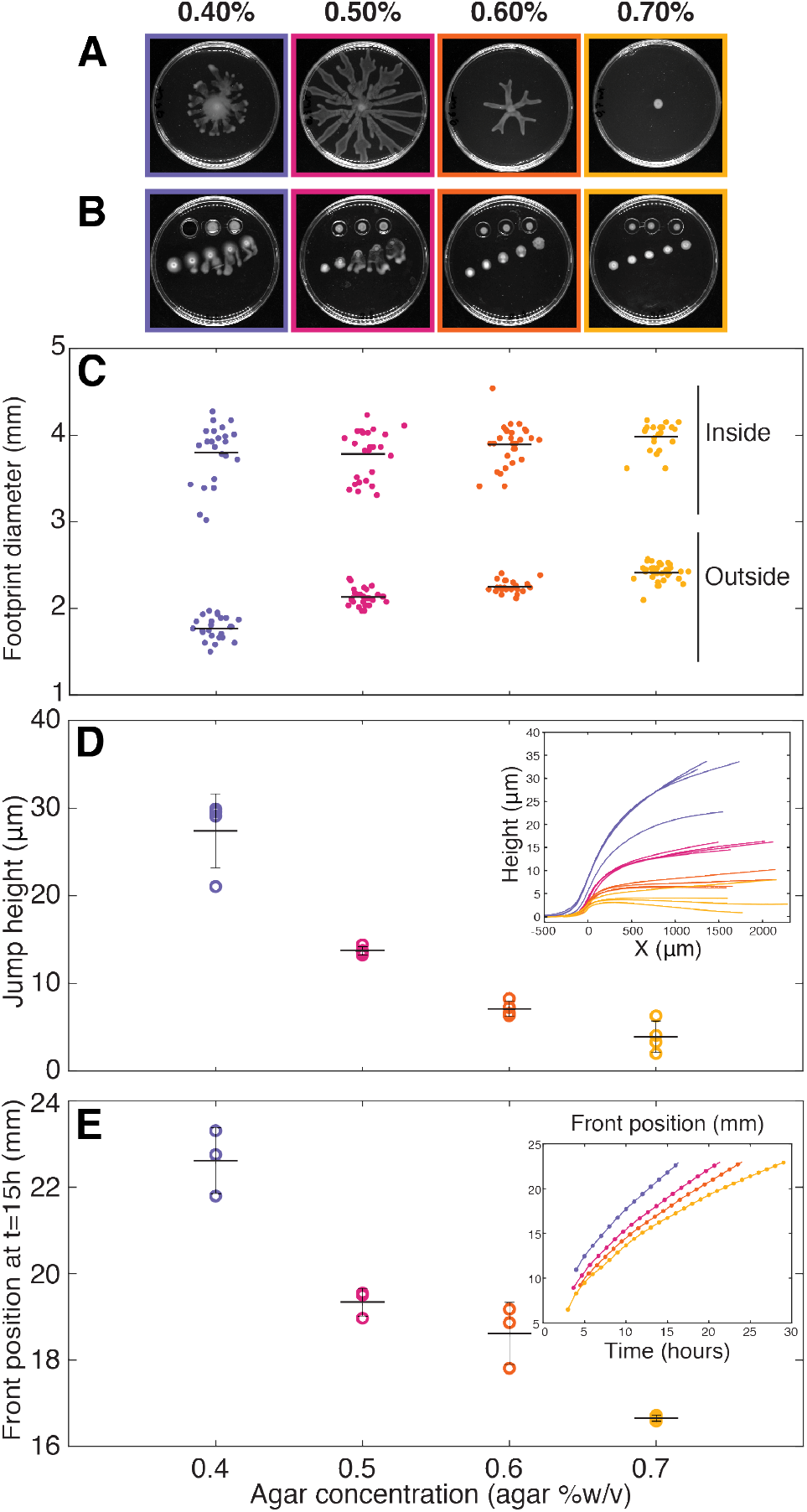
Physical alteration of the gel occurs across a range of agar concentration. A: Snapshots of *P. aeruginosa* swarming colonies, 22 hours after inoculation. Each snapshot is surrounded by a square whose color consistently represents the agar concentration across panels. B: Snapshots of macroscopic rescue experiments, 22 hours after inoculation. (Note one agar patch fell from the two-face Petri dish on the 0.4% plate, just before image acquisition.) C: Footprint diameters for sessile droplet experiments. Droplets were deposited within the rhamnolipid-imbibed area (denoted “Inside”) or outside the swelling front (denoted “Outside”). Data points positions were randomized on the horizontal axis for better visualization. Data points originates from 2 biological replicates. D: Jump height measured 1 mm from the swelling front. Inset: Height profiles obtained by optical interferometry. E: Position of the swelling front 15 hours after inoculation. Data points originates from 3 technical replicates. Inset: Time evolution of the front position, from one replicate. For all panels, horizontal bars are dataset averages and vertical bars are standard deviations.

First, I reproduced the sessile droplet experiment as in Figure 1D. Footprint diameters increased with gel concentration when droplets were deposited on the raw gel surface (outside the rhamnolipid imbibed area), from 1.77 *±* 0.13 mm for 0.4% to 2.41 *±* 0.10 mm for 0.7% (Figure 5C). In contrast, when droplets were deposited inside the rhamnolipids-imbibed area, footprint diameters were overall much greater and independent of agar concentrations. They were about 3.86 (*±*0.27 mm) for all conditions, as if rhamnolipids equalized the wetting properties of the imbibed gels.

Second, I measured the swelling front height profiles generated by rhamnolipid imbibition (Figure 5D). I observed a strong influence of agar concentration: the swelling front jump (measured 1 mm away from the front, see details in Methods) was 27.4 *±* 4.2 µm for 0.4% and gradually decreased for larger agar concentrations (3.9 *±* 1.8 µm for 0.7%). This trend was confirmed by shadowgraphy: the front on a 0.4% gel yielded to a strongly contrasted black line, whereas the front on a 0.7% gel was nearly not visible (data not shown).

Third, I located the position of the swelling front as a function of time, on two-face Petri dishes. A D*rhlA:P*_*BAD*_*rhlAB* colony was seeded on the small aperture on the opposite side, in order to limit the spatial expansion of the rhamnolipid source and to allow for comparison of swelling front advancing speeds across agar concentrations without having to control for the source colony size. The position of the swelling front followed a power law *r* = *t*_*a*_ *with a rang*ing between 0.4 and and 0.6 (Figure 5D, inset), in agreement with the Lucas-Washburn equation for imbibition processes (de Azevedo *et al*., 2008; Cai *et al*., 2021). However, a reproducible estimation of the exponent was difficult, considering the day-to-day variability of agar gel preparation previously described (Tremblay & Déziel, 2008; Ha *et al*., 2014; Pearson, 2019). I turned to a simpler and more robust quantification: the position of the front 15 hours after inoculation. Using this measure, I found that the swelling front advanced significantly faster in low concentration agar gels compared to high concentration agar gels (Figure 5E). These results shed a new light to the well-established fact that swarming motility is strongly dependent on gel hardness (Tremblay & Déziel, 2008; Kamatkar & Shrout, 2011), and demonstrate that swarming colony morphogenesis correlates with the long distance physical alteration of the gel, across a range of agar concentrations, via secretion of rhamnolipids.

## Discussion

Self-produced surfactants play an critical role in a many bacterial genera. For example, the swarming of *Rhizobium etli* (Daniels *et al*., 2006), *Serratia marcescens* (Ang *et al*., 2001), *Bacillus subtilis* (Angelini *et al*., 2009), and *P. aeroginosa* (Caiazza *et al*., 2005) were shown to be linked to the presence of biosurfactants. Here, I documented that rhamnolipids, secrected by *P. aeruginosa*, physically modify the agar gel by inducing a gel swelling. The swelling front displays a maximum angle of 3º that is sufficient to make the edge visible to the naked eye and by shadowgraphy. Previous work had already noted a change of the surface visual aspect. Those studies used terms like “zone of fluid”, “zone of liquid” (Caiazza *et al*., 2005), or “ring of biosurfactants” (Xavier *et al*., 2011) to describe this modified surface state, but my findings show those views were incomplete. Further investigation will be necessary to understand the swelling mechanism, which might involve complex interplay between gel pores, osmolarity, hydrophobic interactions, etc., and might control the spatial distribution of rhamnolipid congeners (Tremblay *et al*., 2007), and goes beyond the scope of this study. Yet, since bacterial colony spreading is controlled by osmotic influx of water from the hydrogel into the colony. Rhamnolipids could possibly be playing a role in this process by causing local swelling in the gel. Investigating this further could be a promising area of research.

Additionally, I demonstrated that transport of rhamnolipids is not restricted to the surface of the agar gel. On the contrary, rhamnolipids imbibe the whole thickness of the gel. A direct consequence of the transport in volume is that spreading speed depends on the local section of the gel, with faster spreading on thinner gels. The modulation of the swelling front traveling speed is likely to be conflated with resource availability: a thinner gel contains less nutrient, yields to a smaller colony, and is expected to produce to a slower swelling front. Experimental data goes in the opposite direction: thinner gels yields to faster, not slower, swelling front. Interestingly, colonies do not spread faster on thinner gel. This highlights the complex interplay between resources availability, rhamnolipids spatial distribution, and swarming motility (Xavier *et al*., 2011; Boyle *et al*., 2015), and the importance of spatial structure in stabilization of cooperative behaviors (Monaco *et al*., 2022; Deforet *et al*., 2019).

The swelling front advancing speed, typically 1 mm/h, is substantially lower than the spreading dynamics of surfactants on liquid films (typically 1 mm/s (Matar & Troian, 1999)). It resembles precursor front in fluid imbibition dynamics of porous materials, which can be interpreted as non-Fickian transport in a Darcy flow (de Azevedo *et al*., 2008; Cai *et al*., 2021). Moreover, the deformation of the agar gel is likely to be coupled to the effective diffusivity of rhamnolipids through the gel. Since rhamnolipids are capable to reach the opposite surface, one could assume, in first approximation, a uniform concentration profile across the gel thickness. Yet, agar gel is a physically-crosslinked hydrogel, and the physico-chemistry of its interaction with surfactants could lead to a non-uniform profile of concentration (Banaha *et al*., 2009; Boral *et al*., 2010). Colony branching in two-face rescue experiments confirms that bulk gel imbibition generates a lateral gradient of surfactant concentration that could potentially interact with the spreading colony itself.

Alteration of the biotic or abiotic environment by living organisms have the potential to modulate motility and dispersal mechanics. Secreted metabolites can act as chemoattractants (Roussos *et al*., 2011; Colin & Sourjik, 2017), trigger quorum sensing (Waters *et al*., 2005), participate in self-organization in stigmergic behavior (Theraulaz & Bonabeau, 1999; D’alessandro *et al*., 2021), and control single-cell motility (Wershof *et al*., 2019). These mechanisms rely on release of diffusible molecules, or local deposition of non-diffusible signals. Here, I evidenced a mechanism where bacteria alter their physical world on a lengthscale substantially larger that their own size. I confirmed this effect holds true across a range of agar concentration. It would be interesting to test other biogel materials (Morin & Déziel, 2021), especially those with biomedical implications, such as mucus (Yeung *et al*., 2012; Rossy *et al*., 2022). Moreover, rhamnolipids are shown to be involved in many killing processes (towards other bacteria (Bharali *et al*., 2013), fungus (Soltani Dashtbozorg *et al*., 2016), immune cells (Jensen *et al*., 2007), or higher organisms (Zaborin *et al*., 2009; Silva *et al*., 2015)): understanding how this killing agent is transported through complex media is of critical importance.

This complex interplay between surfactants secretion, biogel properties alteration and bacterial motility is a unique example of the capacity of bacteria to change the mechanical properties of the world around them, and ultimately, to interact with their distant peers (Estrela *et al*., 2019).

## Acknowledgments

I thank members of the Laboratoire Jean Perrin for fruitful discussions. I am grateful to Françoise Brochard-Wyart for pointing me to the reference (de Azevedo *et al*., 2008), and to Eric Deziel for providing the *rhlB*^*-*^ *and rhlC*^*-*^ *s*trains used in this study. This project has received financial support from the CNRS through the MITI interdisciplinary program, from the Alliance Sorbonne Université through the Emergence program, and from the ANR (ANR-21-CE30-0025).

## Supplementary figures

**Figure S1:**
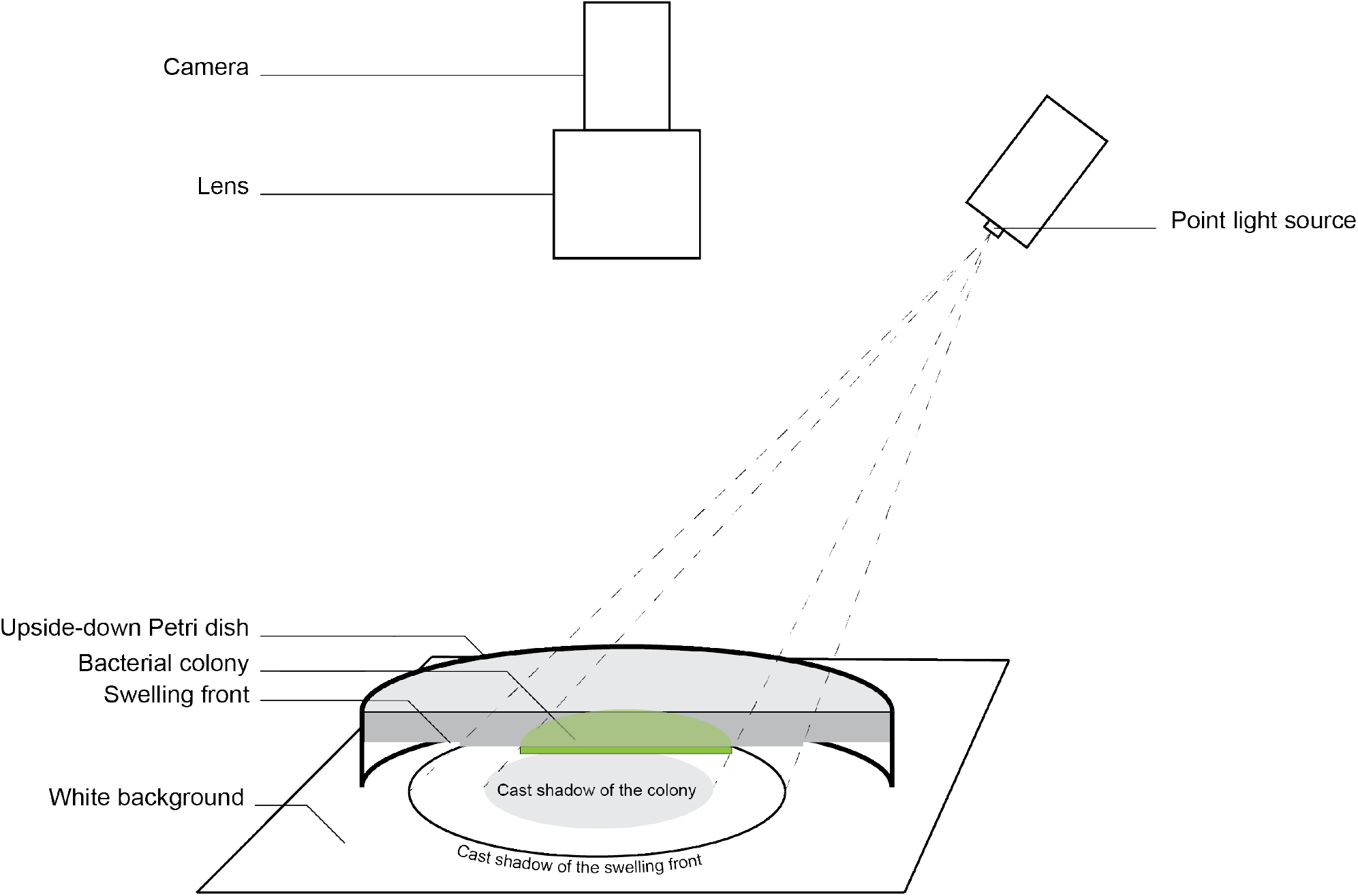
Schematic of the shadowgraphy principle: the light coming from a point source illuminates the colony, but its shadow is cast onto a white background. The camera is focalized on the white background to record the shadow, not on the surface of the gel. For clarity, only half of the Petri dish is displayed.

**Figure S2:**
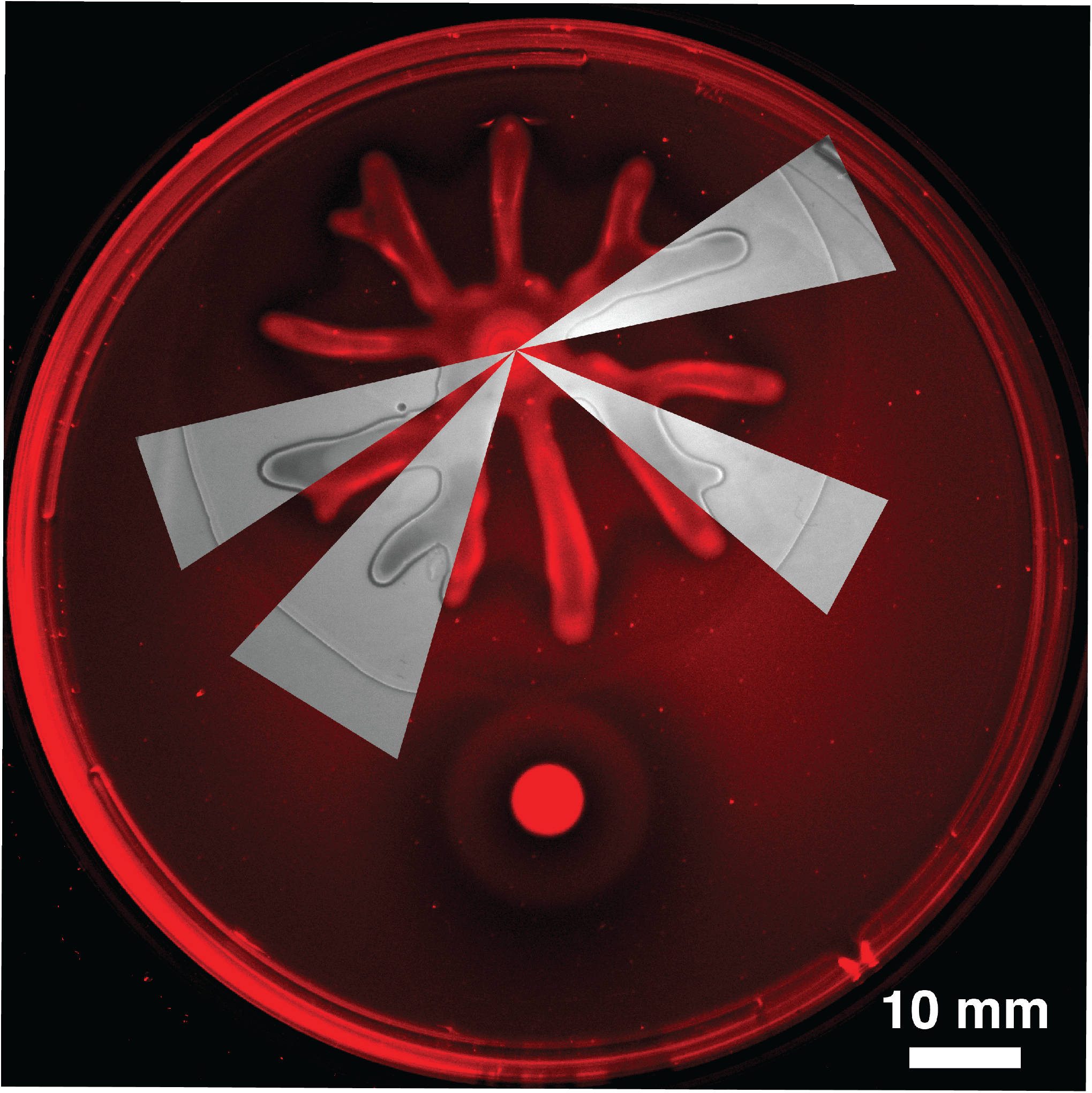
Combined snapshots from shadowgraphy and Nile Red fluorescence imaging. The swelling front revealed from shadowgraphy coincides with a drop of fluorescence that corresponds to a change of solvent polarity.

**Figure S3:**
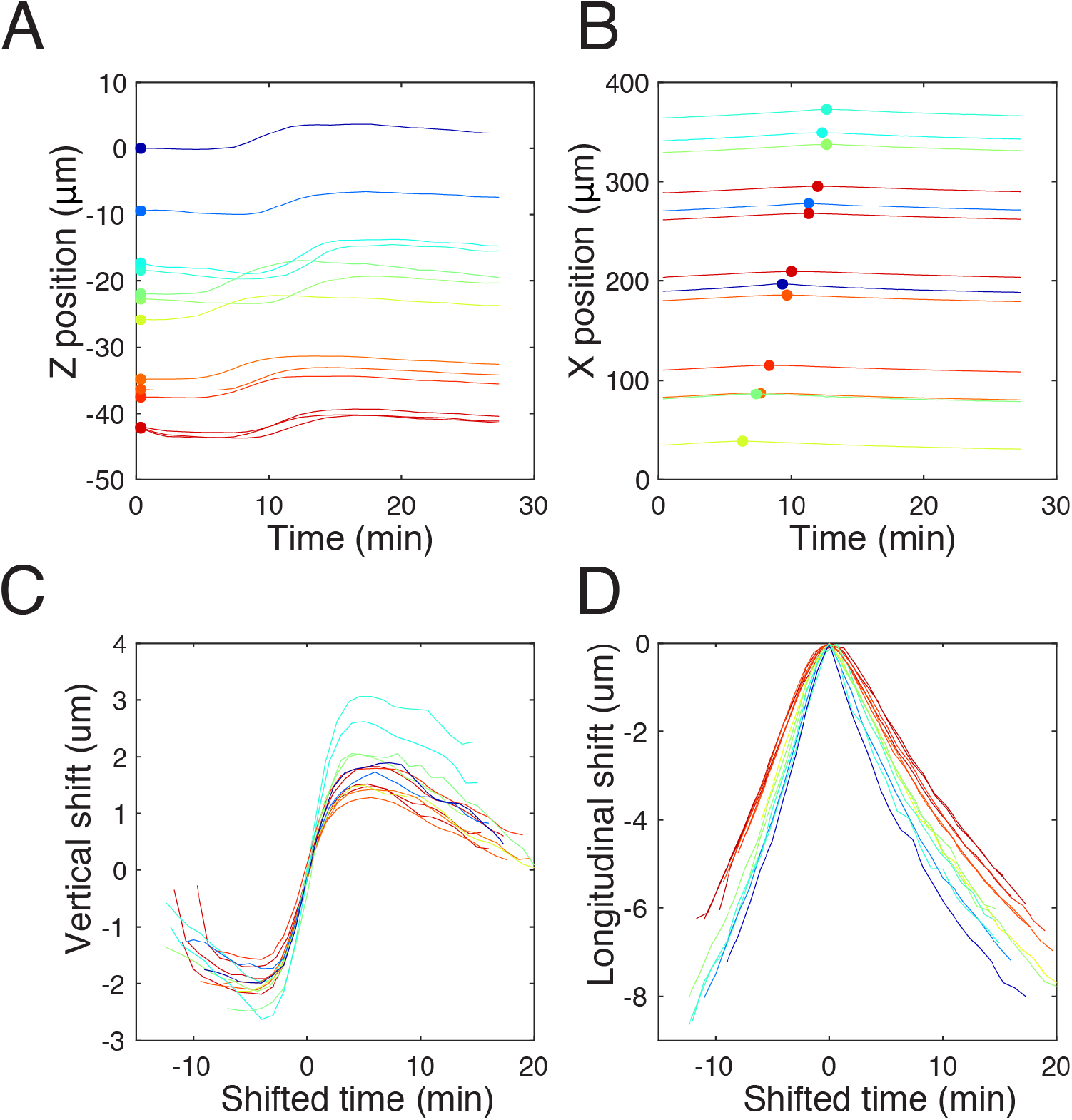
The swelling front is characterized by vertical shift and longitudinal dilation. A: Absolute position *Z*(*t*) for all beads (N=13) detected and tracked during this acquisition. Color in all panels codes for initial depth in the gel. B: Absolute position *X*(*t*) for all beads. The maximal *X* position for each bead is depicted with a dot. The (*X*_*max*_, *t*_*max*_) coordinates of these dots reveals the propagation speed of the swelling front. C: Relative vertical positions (*Z*(*t*) *- Z*(*t*_*max*_)) for all beads, as a function of relative time (*t - t*_*max*_). The swelling amplitude is independent on the initial depth. D: Relative longitudinal positions (*X*(*t*) *- X*_*max*_), as a function of relative time (*t - t*_*max*_). Deeper beads move slower in the longitudinal direction. For the sake of clarity, Figure 1F only represents a subset of the data.

**Figure S4:**
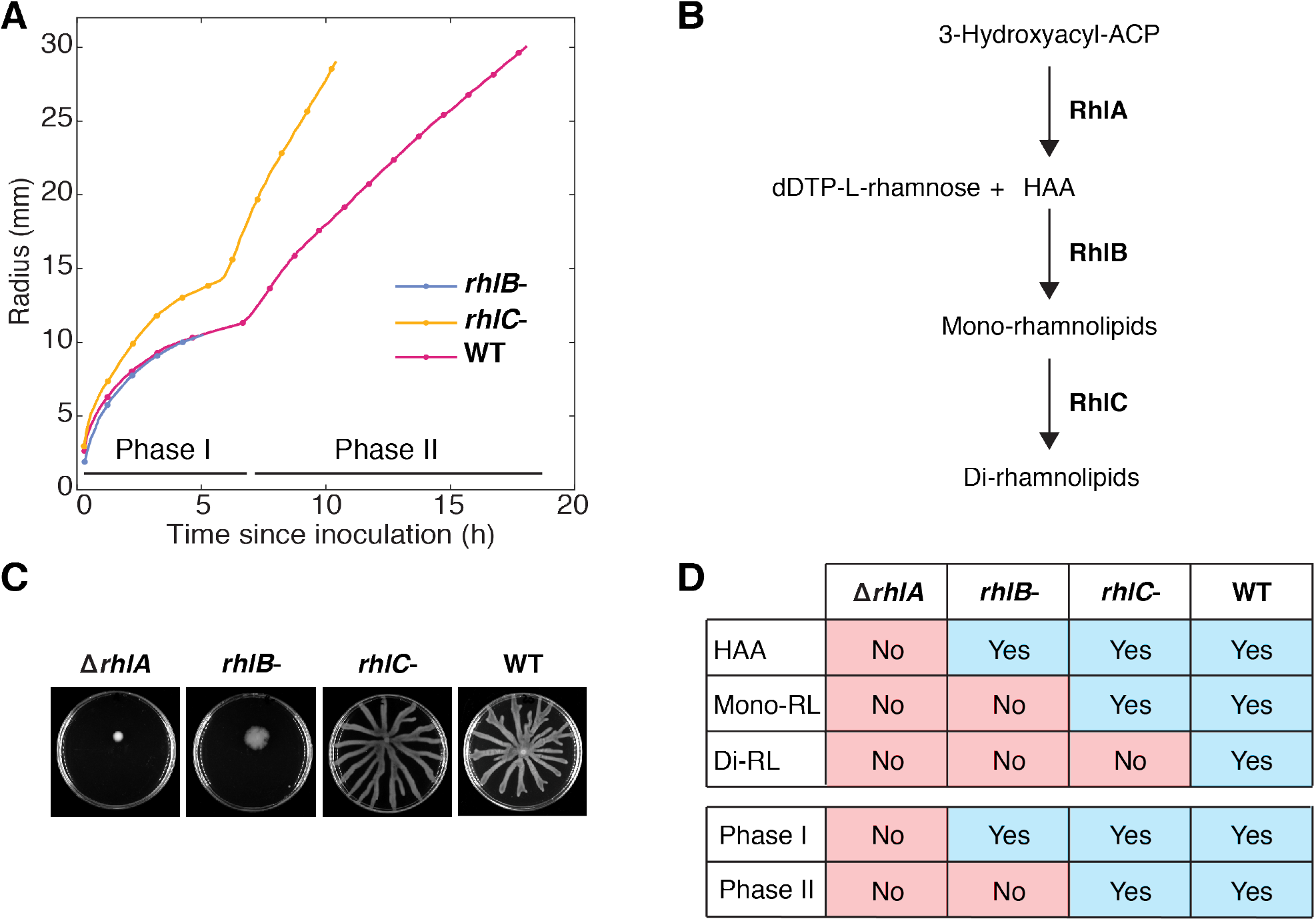
A: Position of the swelling front with respect to time, for *rhlB*^*-*^, *rhlC*^*-*^, *an*d WT colonies. Biosynthesis pathway of rhamnolipids, in three steps, involving RhlA, RhlB, and RhlC proteins. C: Swarming colony for the four strains, 24 hours after inoculation. D: Summary table comparing rhamnolipid secretion capabilities of the four tested strains, with the presence of Phase I and Phase II in curves from panel A.

**Figure S5:**
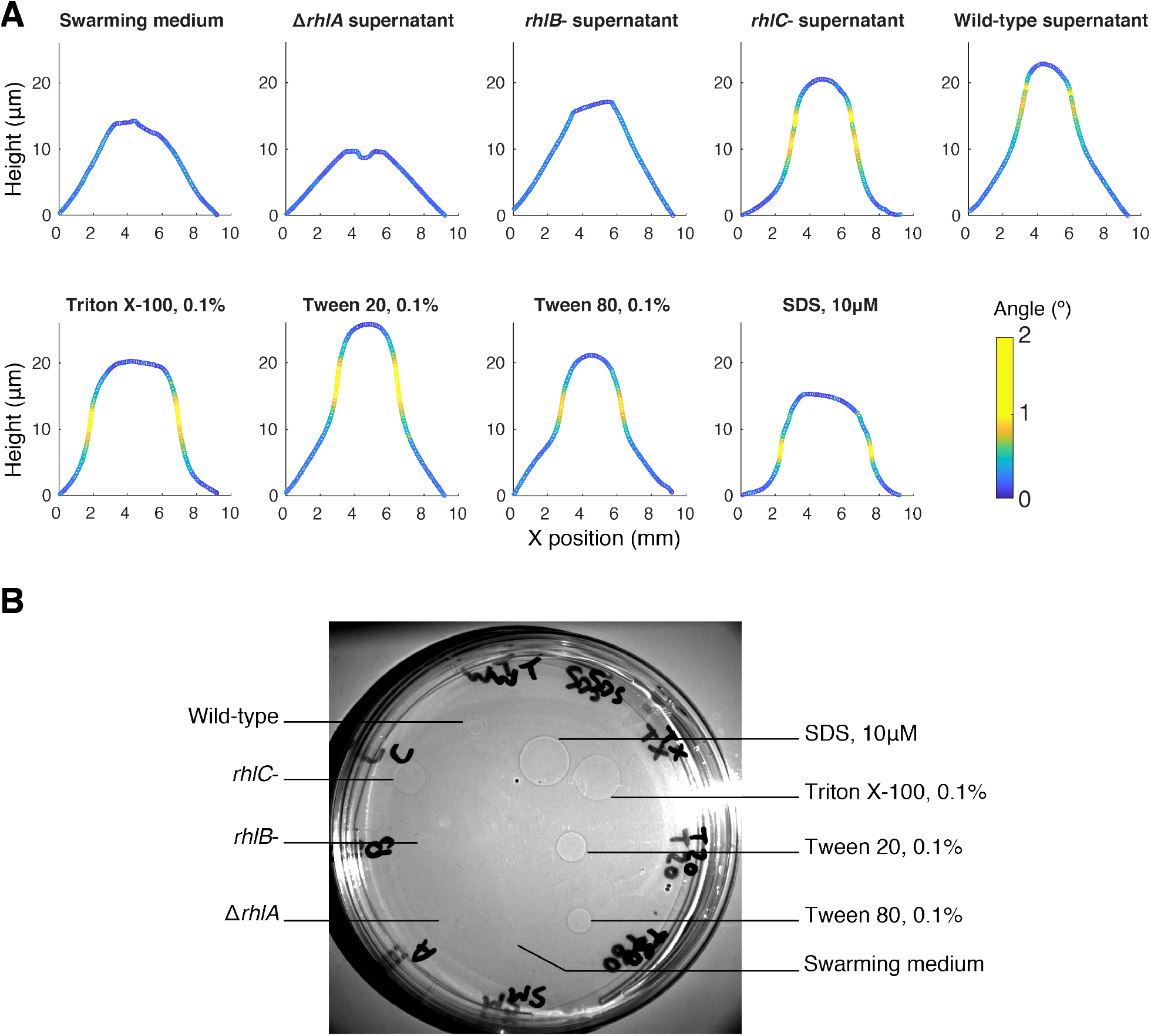
A: Profiles of the gel surface obtained with an optical profilometer, 5 minutes after deposition of a 1 µL droplet. Nine samples were tested: Swarming medium (i.e. media use for swarming plates, minus agar), supernatants of D*rhlA, rhlB*^*-*^, *rhlC*^*-*^, *an*d wild-type culture in swarming media, and four synthetic surfactants: Triton X-100, Tween 20, Tween 80, and SDS. Note the vertical scale is approximately enlarged 1000-fold compared to the horizontal scale: the gel remains very flat in all conditions, except in regions corresponding to the swelling fronts, highlighted here in yellow, where the curve slope is greater than 1º. B: Shadowgraph of an agar plate, taken 20 min after deposition of 2 µL droplets of the 9 tested samples. Only the four synthetic surfactants and the supernatants of wild-type and *rhlC*^*-*^ *cultures yi*eld to a swelling front. Swarming medium, and supernatants of D*rhlA* and *rhlB*^*-*^ *do not indu*ce a swelling front.

**Figure S6:**
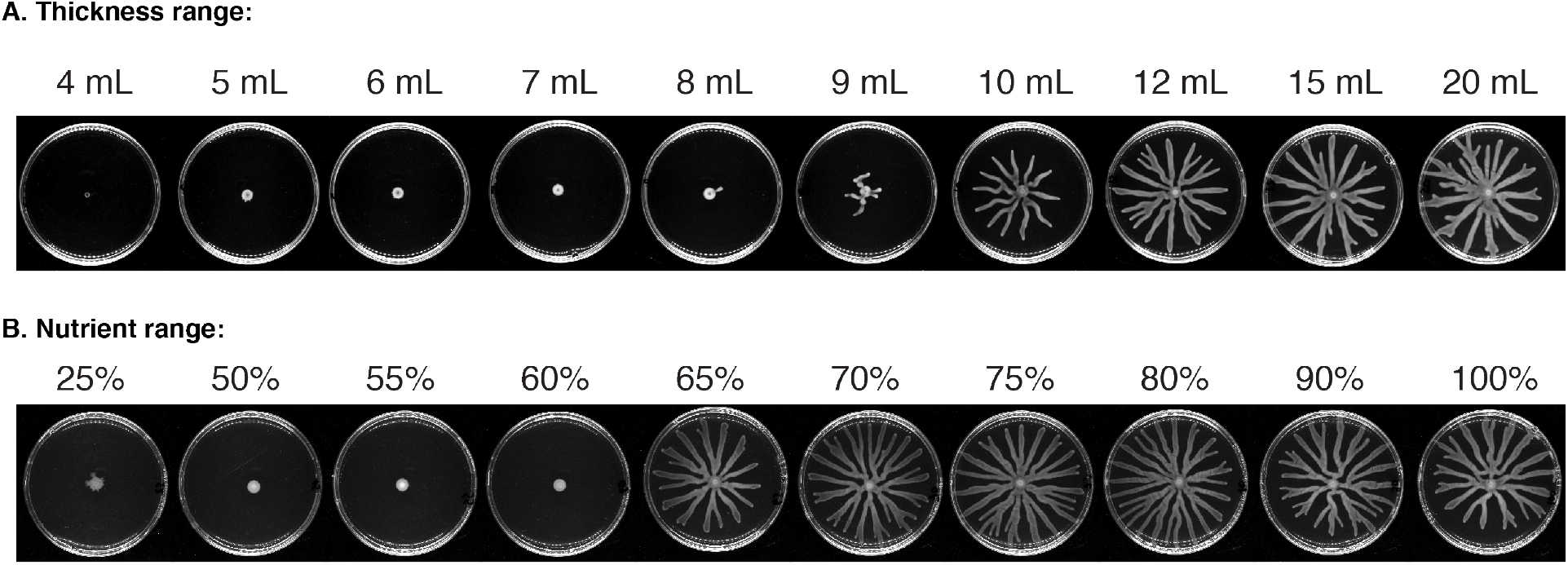
A: Snapshots of swarming colonies grown on gels of various thickness, expressed as volume of agar gel in the plate. 20 mL is the nominal volume, and corresponds to a thickness of 3.5 mm. B: Snapshot of swarming colonies grown on a gel containing various concentration of nutrients (casa-aminoacids). The nominal concentration is 12.5 g/L. Concentrations are expressed as a percentage of this nominal concentration. Note the change of final colony size is abrupt between 60% and 65% concentrations of nutrients, while the change is more gradual between 8 mL and 12 mL volumes. All snapshots were taken 22 hours after inoculation.

**Figure S7:**
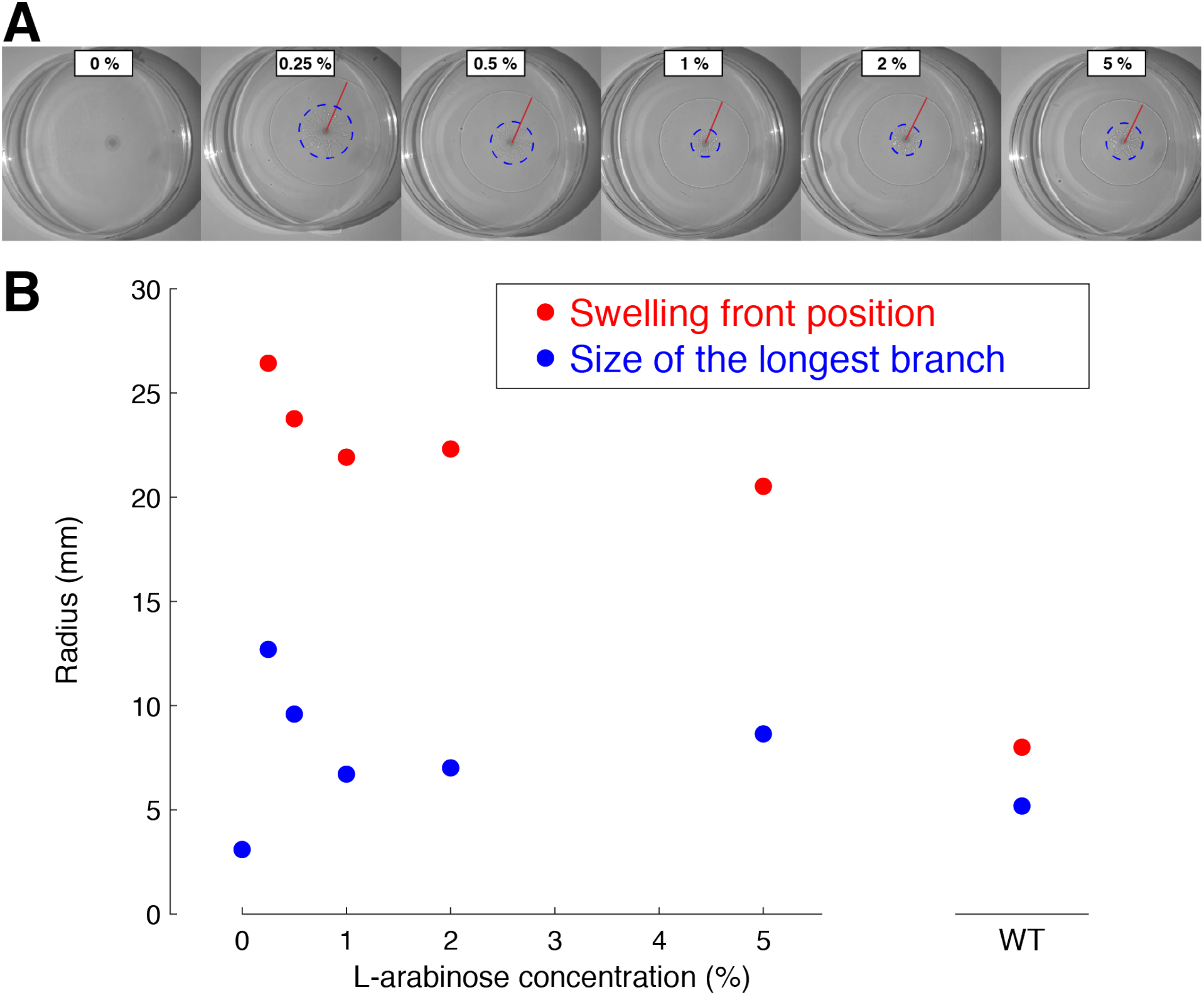
A: Shadowgraphy snapshots, taken 8 hours after inoculation, of D*rhlA:P*_*BAD*_*rhlAB* colonies growing on agar gel plates supplemented with varying concentrations of L-arabinose. The distance between the inoculation point and the swelling front is highlighted in red, the longest branch of the colony sets the radius of the dashed circle. B: Position of the swelling front (red dots) and colony sizes (blue dots) are reported with respect to L-arabinose concentration. For comparison, swelling front position and colony size for a wild-type colony 8 hours after inoculation are also reported. The concentration of 1% of L-arabinose, chosen for all rescue experiments, corresponds to the smallest colony size while maintaining an abundant production of rhamnolipids (evaluated by the position of the swelling front).

**Figure S8:**
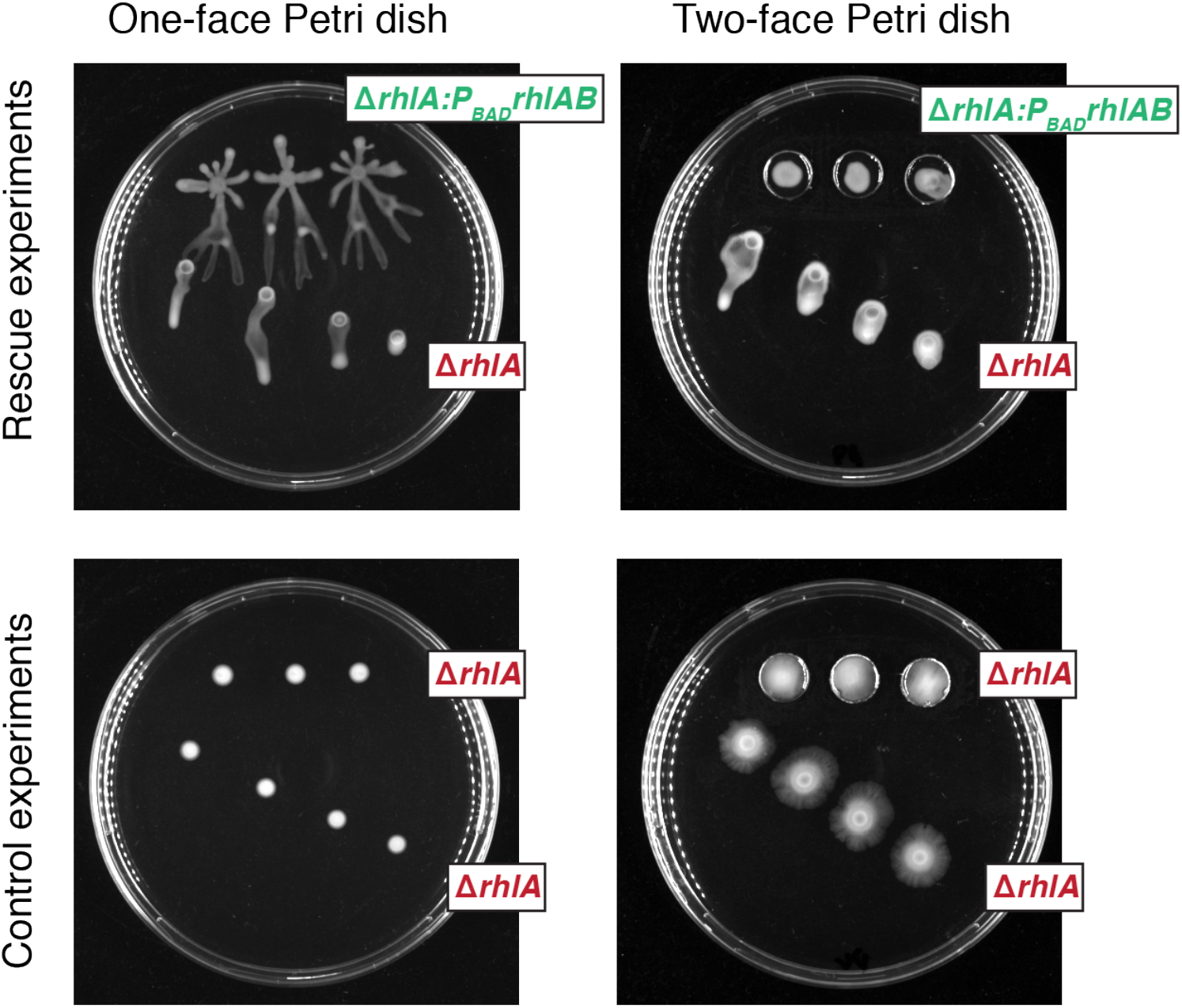
Rescue experiments (top) and control experiments (bottom). Left: on one-face Petri dish. Right: on two-face Petri dish. D*rhlA* colonies are unable to swarm by themselves. They are rescued using rhamnolipids provided by D*rhlA:P*_*BAD*_*rhlAB* colonies. When facing another D*rhlA* colony, they are not rescued. Snapshots were taken 15 hours after inoculation for one-face Petri dish, 36 hours after inoculation for two-face Petri dish.

## Supplementary Movies

**Movie S1:**
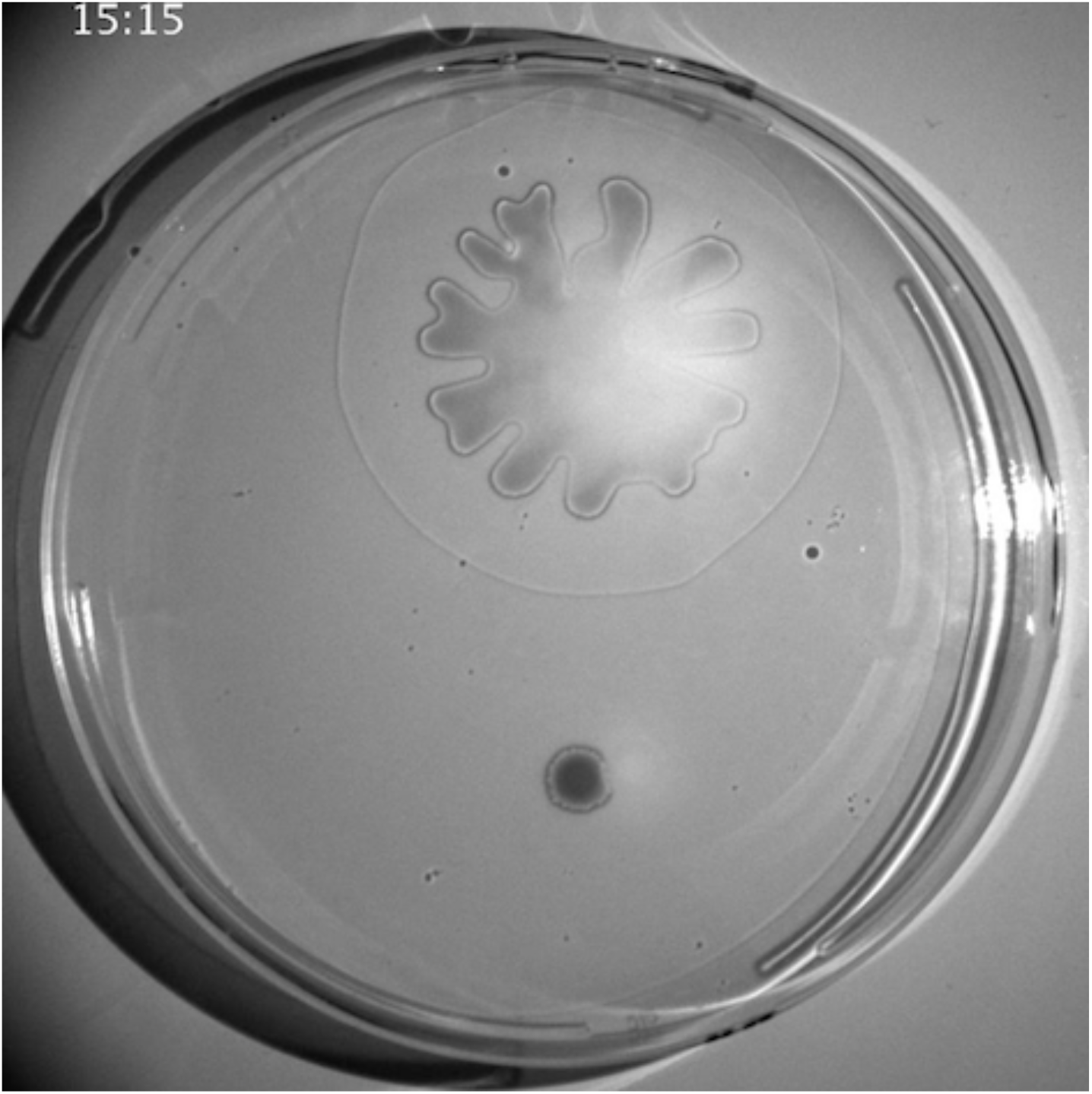
Wild-type vs D*rhlA* swarming colonies captured in shadowgraphy

**Movie S2:**
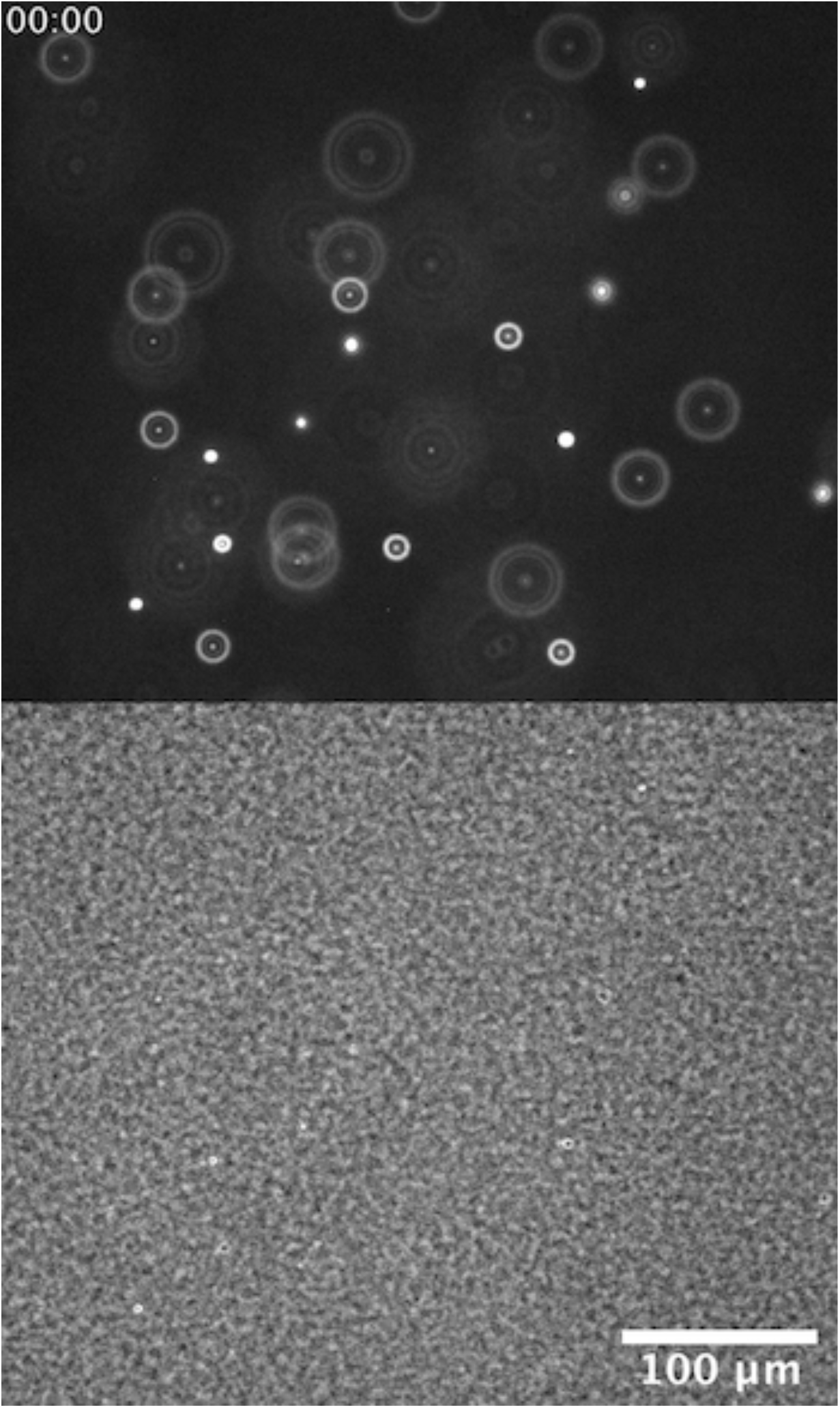
Timelapse video of 1 µm fluorescent beads embedded into the agar gel. Top: fluorescence image. Bottom: phase contrast image. The swelling front passing in the field of view (from left to right) is visible as a white halo in the phase contrast image, from t=4 min to t=15 min.

**Movie S3:**
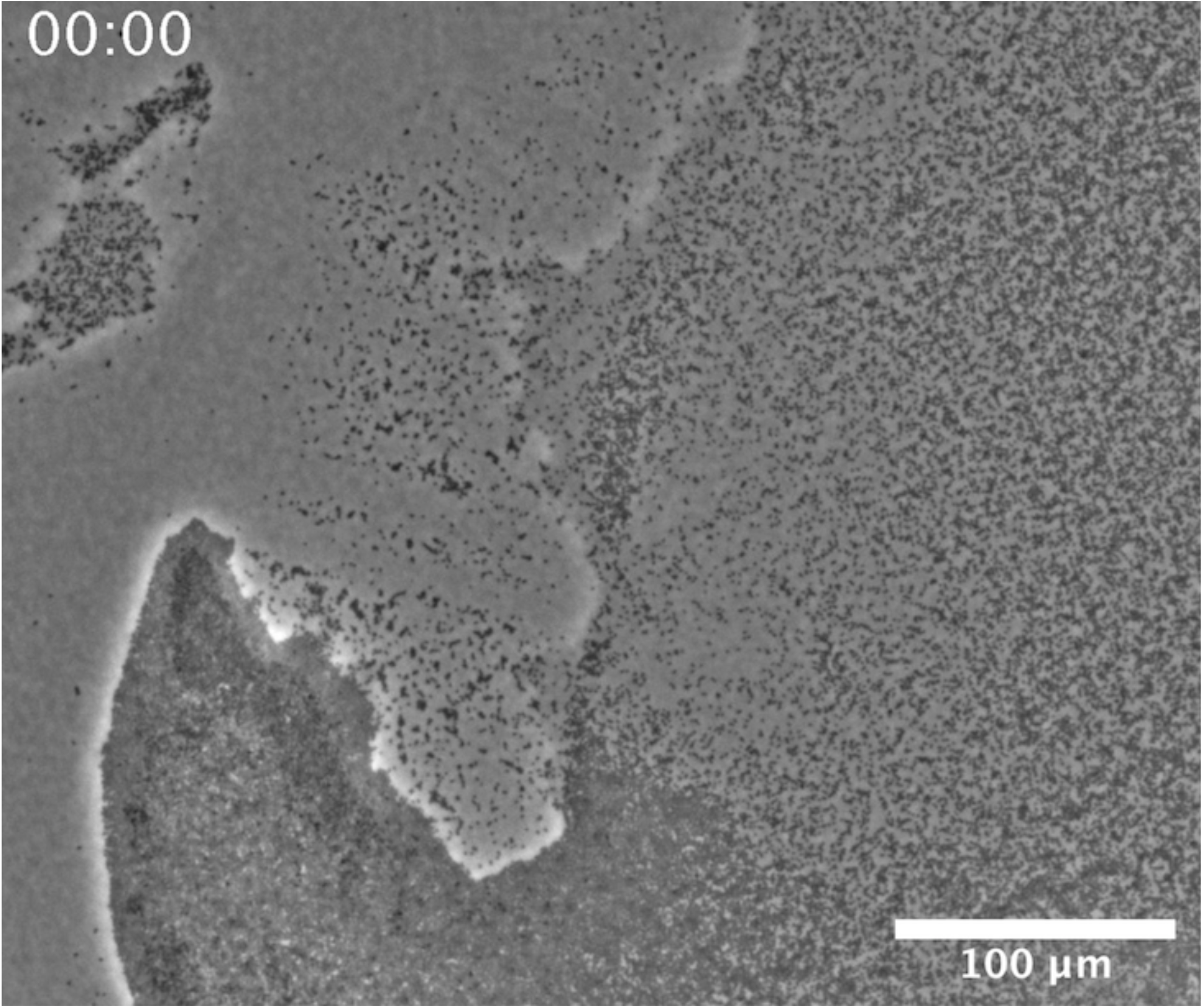
Timelapse video of D*rhlA* cells motility on agar gel. The swelling front passes through the field of view at t=20 min, from the left side to the right side.

**Movie S4:**
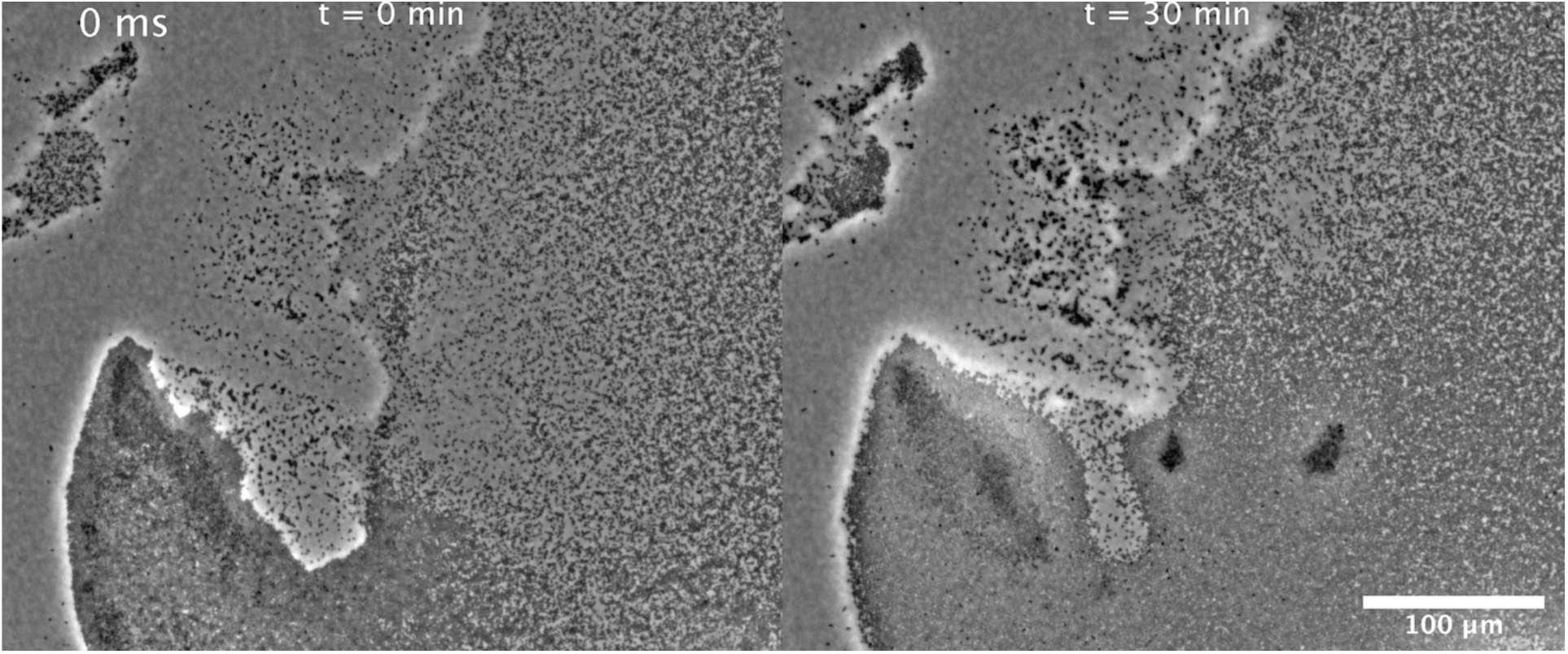
One second long videos of D*rhlA* cells motility on agar gel. Left: t=0 min, before passage of the swelling front. Right: t=30 min, after passage of the swelling front.

**Movie S5:**
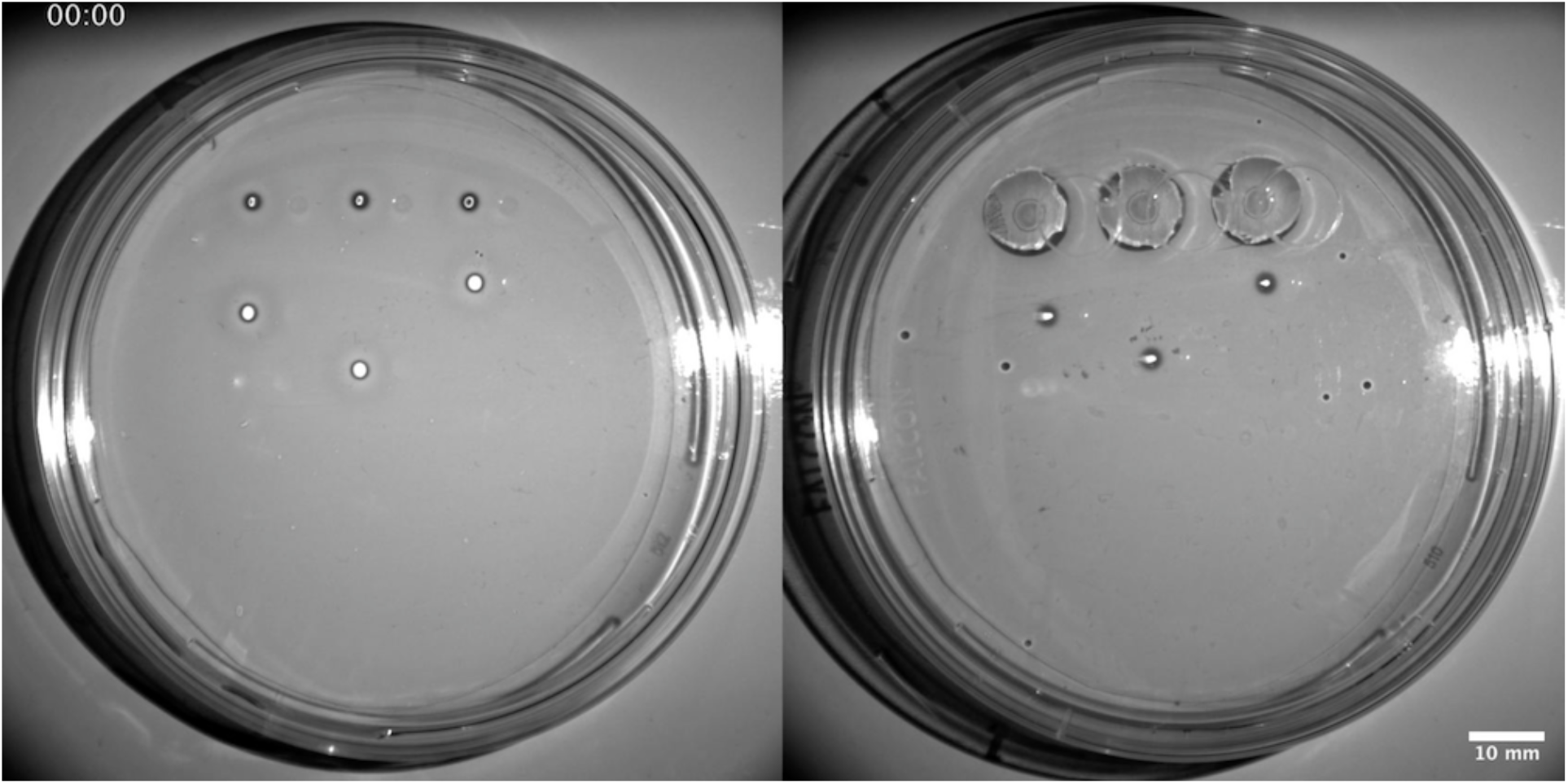
Left: Rescue experiment on a regular one-face swarming plate. Right: Rescue experiment on a two-face swarming plate. In both cases, three D*rhlA* colonies, unable to swarm by themselves, are rescued using rhamnolipids provided by three D*rhlA:P*_*BAD*_*rhlAB* colonies grown at a distance (ranging from 7 to 23 mm). On the two-face plate, D*rhlA:P*_*BAD*_*rhlAB* colonies are grown on the second face, within holes. On the one-face plate, D*rhlA:P*_*BAD*_*rhlAB* colonies are grown on the same face as the D*rhlA* colonies. Note that the left and center D*rhlA* colonies start swarming before the D*rhlA:P*_*BAD*_*rhlAB* colonies reached them (respectively 90 minutes and 140 minutes). Full movie duration is 20 hours.

